# Hierarchical organization in sparse gene regulatory networks shapes structural coherence and emergent regulatory coordination

**DOI:** 10.64898/2026.02.04.703680

**Authors:** Pradyumna Harlapur, Rahul Jagadeesan, Andre S. Ribeiro, Claus Kadelka, Mohit Kumar Jolly

## Abstract

A fundamental challenge in systems biology is understanding how regulatory coordination is maintained within extremely sparse gene regulatory networks (GRNs). While traditional unsigned network metrics can quantify the overall magnitude of influence a specific node or edge exerts over the network, they cannot predict the functional direction and consistency of these regulatory changes. To address this gap, we introduce the coherence matrix. This topology-based framework for signed, directed GRNs computes a strictly bounded, inter-pretable measure of directional regulatory coordination between gene pairs. Using synthetic network ensembles, we demonstrate that architectures with low intrinsic structural coherence suffer severe density-driven scaling penalties. Such structures cannot scale without requiring biologically unrealistic edge densities. Applying this framework to six whole-organism GRNs reveals how biological topologies overcome these limitations. Whole-organism GRNs exhibit lower overall coherence and more fragmented structures than expected, avoiding rigid global co-regulation to maximize regulatory degrees of freedom. Furthermore, these systems resolve the scaling penalties of complex, low-coherence architectures by restricting dense regulatory crosstalk to a compact intermediate core. Modeling synthetic hierarchical networks confirms that this architecture—a compact core within a globally feed-forward scaffold—neutralizes scaling penalties by minimizing nodes requiring dense connectivity. This allows GRNs to efficiently route decisions despite extreme sparsity. Finally, transcriptomic validation confirms the coherence matrix accurately predicts empirical co-regulation directionality. By moving beyond unsigned connectivity, the structural coherence framework provides a quantitative tool to map how the intrinsic behavior of localized interactions scales into macroscopic, system-wide coordination.

## 1 Introduction

Gene regulatory networks (GRNs) enable cells to generate precise and context-dependent gene expression programs in response to internal and environmental cues. GRNs function as information-processing systems that maintain coordinated behavior despite stochastic noise, genetic perturbations, and the inherent sparsity of regulatory connections [1, 2, 3, 4]. Here, regulatory coordination refers to the consistency of regulatory influence transmitted between genes through both direct and indirect pathways.

A central challenge in systems biology is therefore to understand how large-scale regulatory coordination emerges from the local topology of directed, signed interactions. Over the past two decades, GRNs have been extensively characterized using graph-theoretic descriptors such as degree distributions [5, 6, 7], modularity [8, 9, 10, 11], hierarchical organization [12, 13, 14], and the enrichment of specific network motifs [15, 8, 16, 17, 18, 19, 20]. These approaches have revealed important organizational principles, including scale-free connectivity [5, 6], modular structure [8, 9], and feedback-dominated cores [13]. However, connecting these structural features to the consistency of regulatory influence across the network remains unresolved. In particular, coordination in GRNs is not determined solely by local connectivity or direct interactions, but rather emerges from the collective effects of many interacting regulatory paths.

Standard adjacency-based representations capture immediate regulatory relationships but fail to account for the cumulative logic of signal propagation through a network. In GRNs, regulatory influence is transmitted along signed regulatory paths of varying lengths that may reinforce or counteract one another depending on the signs of the constituent edges [17]. Because GRNs are directed and signed, the functional effect of an indirect path depends on the balance of activating and inhibitory interactions along that path. While traditional metrics that ignore edge signs can possibly identify highly central nodes, they fail to predict the direction of that influence and, consequently, cannot determine how perturbing a specific node will alter the downstream state of the network. To address this gap, we introduce the coherence matrix, a network-topology-based framework that integrates edge signs to quantify the directional consistency of regulatory influence across all pairs of genes. Unlike established signed network metrics [21, 22, 23, 24, 25], which typically conflate balanced regulatory conflict with a complete absence of connectivity by assigning both a value of zero, the coherence matrix explicitly isolates topological disconnection by propagating undefined values for unlinked pairs. Furthermore, existing measures generally yield unnormalized or unbounded values that obscure the actual proportion of activating versus inhibitory signals. These combined limitations, conflating disconnection with conflict and lacking standardized bounds, make it difficult to interpret the relative magnitude of regulatory influence and to compare topologies across different networks. By normalizing the net signed influence against the total path count, the coherence matrix produces a strictly bounded, interpretable measure of true regulatory balance. These structural advantages make the framework uniquely suited for analyzing biological GRNs, which are inherently sparse and hence dominated by unconnected gene pairs.

In this study, we apply the coherence framework to investigate how local regulatory logic scales to maintain global coordination under the severe biological constraint of network sparsity. We first utilize synthetic networks to demonstrate that a circuit’s intrinsic structural coherence and topological class dictate its resilience to sparsification. Extending this framework to six whole-organism GRNs, we reveal how biological systems utilize network topology to overcome intrinsic scaling limitations. Evaluating the distribution of structural coherence demonstrates how these networks naturally partition into independent, mutually activating functional modules, or “teams” [22, 26]. Furthermore, by integrating indirect paths to track how local circuits transition through higher-order interactions, we show that complex, low-coherence architectures persist despite their density-dependent scaling penalties because they are concentrated within a small, densely connected intermediate core. This compact core centralizes regulatory crosstalk, routing and maintaining regulatory coordination across these diverse functional modules and teams despite the stringent biological constraint of extreme global sparsity. By modeling synthetic hierarchical networks, we confirm that this macroscopic topology explicitly neutralizes density-driven scaling penalties, allowing the network to successfully maintain cohesive regulatory teams under extreme sparsity, irrespective of their baseline structural coherence. Finally, by validating our topological predictions against empirical transcriptomic data (*E. coli* iModulons [27]), we demonstrate that the coherence matrix can, to a large extent, accurately predict the direction of co-regulation, specifically whether gene pairs contribute in a concordant or discordant manner, within these independent functional modules. Collectively, these results establish structural coherence as a complementary framework for signed GRNs. By moving beyond traditional metrics of node or edge importance to explicitly quantify the direction and sign of regulatory influence, this approach links localized regulatory logic to the emergence of system-wide coordination under extreme biological sparsity.

## 2 Results

### 2.1 Structural Coherence and Team Organization in Signed Regulatory Circuits

We began our analysis at the level of regulatory circuits, which constitute the fundamental building blocks of gene regulatory networks. Circuits such as the toggle switch, toggle triad, and other recurrent architectures have been extensively studied in biological contexts ranging from cell fate determination and differentiation to developmental patterning and stress responses [28, 29, 30, 15, 31, 32, 33, 34, 19, 20]. These circuits implement distinct regulatory logic—such as mutual inhibition [34, 32], positive feedback, or mixed-sign interactions [28, 29]—that drive characteristic dynamical behaviors such as oscillations or multistability. In whole-organism networks, individual genes rarely act in isolation [35]. Instead, small regulatory circuits often serve as coarse-grained abstractions for larger sets of co-expressed genes acting in concert. Because these motifs rely on downstream effectors, evaluating minimal, unscaled circuits fails to capture the structural vulnerabilities that emerge in high-dimensional biological modules. Consequently, systematically evaluating how local regulatory logic scales establishes a critical baseline for determining which topologies can survive biological sparsification, bridging localized interaction structures with the emergence of global coordination.

To characterize these properties in a topology-driven manner, independent of specific dynamical models, we constructed the coherence matrix (*C*) (Figure 1A) (see Methods). This matrix quantifies regulatory alignment by aggregating the balance of positive and negative interactions over all walks of increasing length. Each matrix entry *C*_*ij*_ ∈ [−1, 1], where values near 1 or −1 indicate exclusively activating or inhibitory influences, respectively, and 0 reflects perfectly balanced conflict. Crucially, unlike existing methods for signed, directed interactions [21, 22, 24], our formulation preserves topological disconnection by propagating undefined entries (NaN). This distinction is vital for sparse GRNs, preventing the conflation of absolute topological disconnection with perfectly balanced conflict between activating and inhibitory interactions.

**Figure 1:**
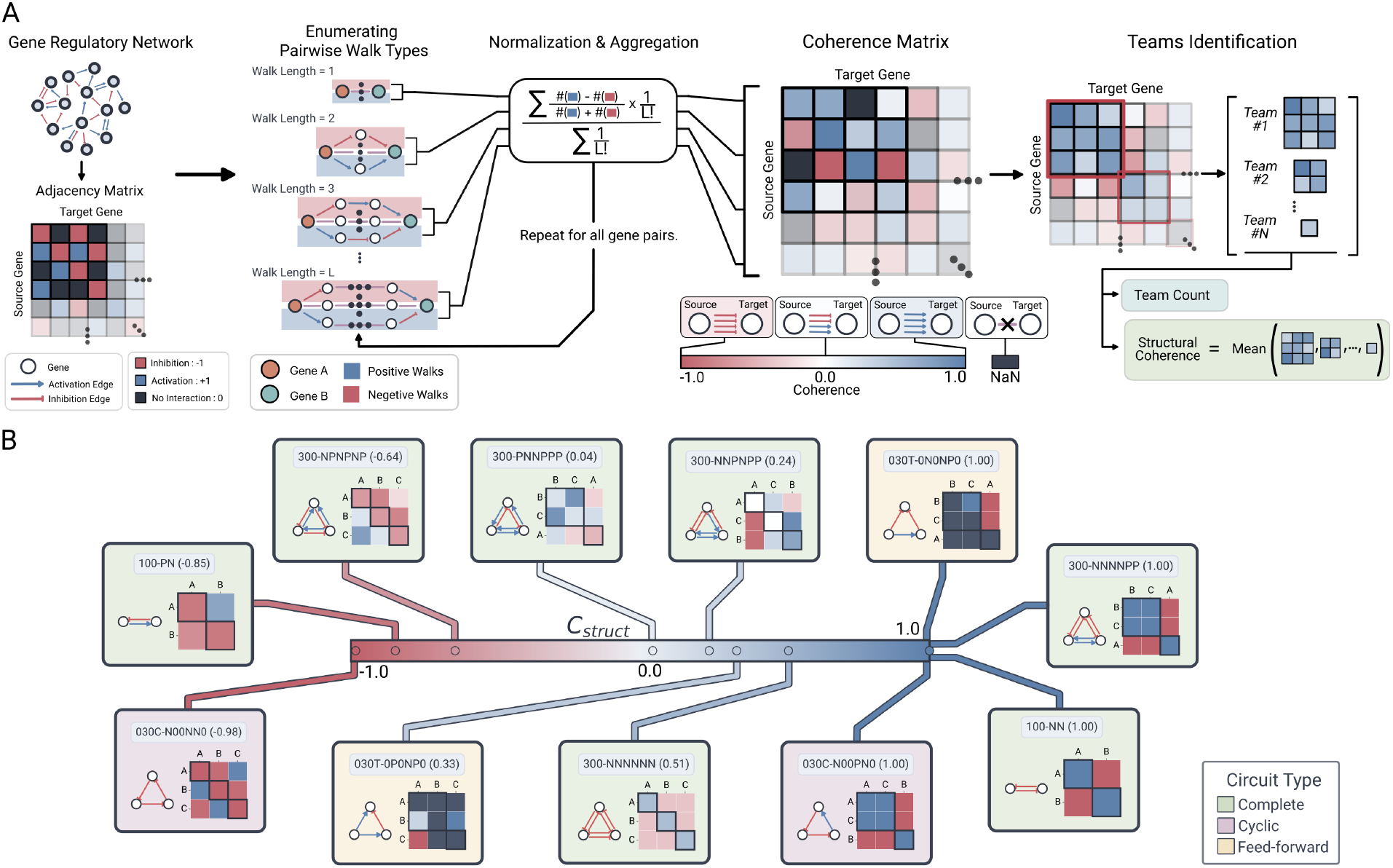
Coherence Matrix Calculation and Structural Coherence Spectrum. (**A**) Schematic pipeline of the network transformation from a signed, directed adjacency matrix **A** into the path-aggregated coherence matrix **C** across successive walk lengths *L* and subsequent post-processing - identifying teams and calculation of structural coherence. (**B**) Representative 2-node and 3-node base circuits (without self-activation) distributed along a continuous spectrum of structural coherence (C_struct_). Visual panels for each sampled circuit depict the MAN topological classification with edge-composition string and its structural coherence value in parenthesis, the directed graph layout (blue arrows: activation; red bar-headed lines: inhibition), and the coherence matrix heatmap. Thick black square borders along the heatmap diagonals highlight the identified teams.

In models explaining biological phenomenon, small circuits frequently serve as coarse-grained abstractions of larger gene sets acting in concert. This widespread co-expression, corroborated by high-throughput transcriptomics, underscores the necessity of coordinated gene sets for executing specific functions [36, 37, 38, 39, 40]. Consistent with prior work, we refer to these mutually activating groups as teams [21, 22]. Within our framework, teams correspond to discrete node subsets strictly devoid of internal inhibitory interactions (all intra-team *C*_*ij*_ *>* 0). To systematically extract these groups, we developed a greedy agglomerative algorithm that identifies maximal subsets of purely activating nodes directly from the coherence matrix (see Methods).

To quantify the overall degree of coordinated interaction within a partitioned network, we defined the metric of *structural coherence* (C_struct_), calculated as the mean coherence value across all within-team interactions (Figure 1A) (see Methods). A structural coherence of 1 indicates that all identified teams consist of nodes that strongly and exclusively activate one another. Conversely, −1 designates a network entirely dominated by negative feedback, fragmenting into isolated, self-inhibiting nodes. Intermediate values reflect varying degrees of mixed regulatory coordination.

### 2.2 Structural Coherence Dictates Robustness of Team Structure Under Sparsification

To determine how regulatory architectures maintain coordination at scale, we evaluated the response of 2-node and 3-node circuits (Figure 1B) to scaling and sparsification. These base circuits span the coherence spectrum and represent three fundamental topological classes: feed-forward (030T), complete 2-node (100), complete 3-node (300), and cyclic (030C) [41] (Figure S1). We expanded each circuit by factors of 10×, 30×, and 50×, replacing individual nodes with groups of nodes equal to the scaling factor. Because team formation requires mutual activation, we generated two parallel sets of networks: Self-Activation (SA) networks, where explicit self-activating loops were added to each base node prior to scaling, and Non-Self-Activation (NS) networks, which lacked these localized loops. This comparison distinguishes coordination driven by direct auto-regulation from that maintained by indirect, network-wide feedback. For each scaled network, we systematically varied global edge density from 10% to 100% in 10% increments. To isolate the effects of progressive sparsification across all these synthetic Erdős–Rényi (ER) network ensembles, we generated 50 independent replicates per configuration using an additive edge-growth approach (see Methods and Supplementary Methods). We then assessed structural divergence from the fully connected (100%) ideal by calculating the absolute error in team count (|ΔTeams|) and structural coherence (|ΔC_struct_|):

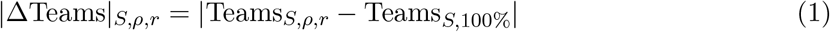

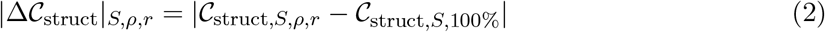

Circuit topological class fundamentally dictated the network’s response to sparsification. Cyclic and complete circuits exhibited progressive structural degradation as edge density decreased (Figure 2A–B, i, iii–iv), whereas feed-forward architectures demonstrated remarkable robustness (Figure 2A-B, ii). The magnitude of this disruption was governed by the base circuit’s absolute structural coherence (|C_struct_|_basal_). When evaluating the absolute error in team count (|ΔTeams|), networks scaled from highly coherent circuits maintained negligible deviation across all density regimes; however, as basal |C_struct_| decreased, sparsification-induced errors in team formation increased significantly (Figure 2A). The absolute error in structural coherence (|ΔC_struct_|) closely followed this pattern, scaling in severity alongside lower absolute basal coherence (Figure 2B, Figure S2). These trends remained consistent across all scaling factors for both SA and NS networks (Figure S3, Figure S4). Furthermore, the shapes of these error trajectories were predictable based on the circuit’s initial |C_struct_|. The most incoherent circuits displayed an immediate, convex sensitivity to sparsification, yielding high error margins even at 90% global density, whereas intermediately coherent circuits exhibited concave trajectories, indicating initial structural resilience prior to eventual regulatory collapse.

**Figure 2:**
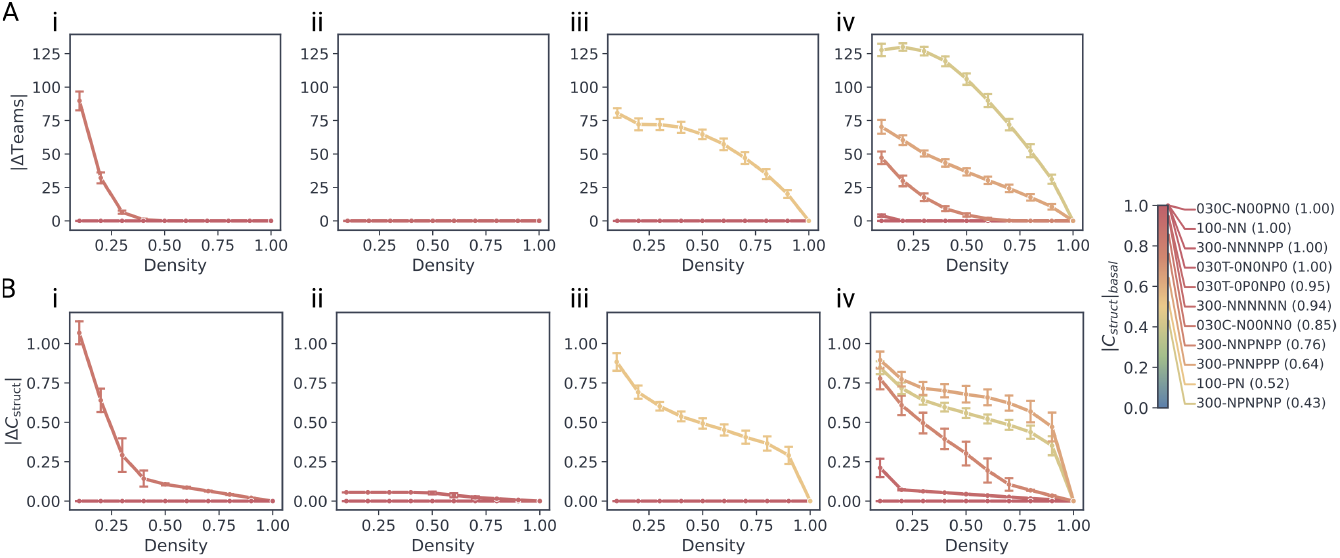
Absolute error trajectories under network sparsification. Line plots tracking the (**A**) absolute error in team count (|ΔTeams|) and (**B**) structural coherence (|ΔC_struct_|) as a function of global edge density (%) for the 50× scaled networks of the self-activation variants of the circuits. Columns segregate results by topological class: (**i**) cyclic, (**ii**) feed-forward, (**iii**) complete 2-node, and (**iv**) complete 3-node architectures. Individual trajectories are colored by the specific base circuit’s structural coherence, and error bands denote the standard deviation across 50 independent replicates per data point.

To quantify net structural susceptibility across the entire sparsification trajectory, we calculated the Cumulative Absolute Error (CAE):

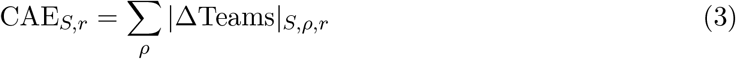

As expected, CAE scaled proportionally with the network scaling factor, demonstrating that larger networks possess a higher capacity for absolute regulatory discordance (Figure 3A,

**Figure 3:**
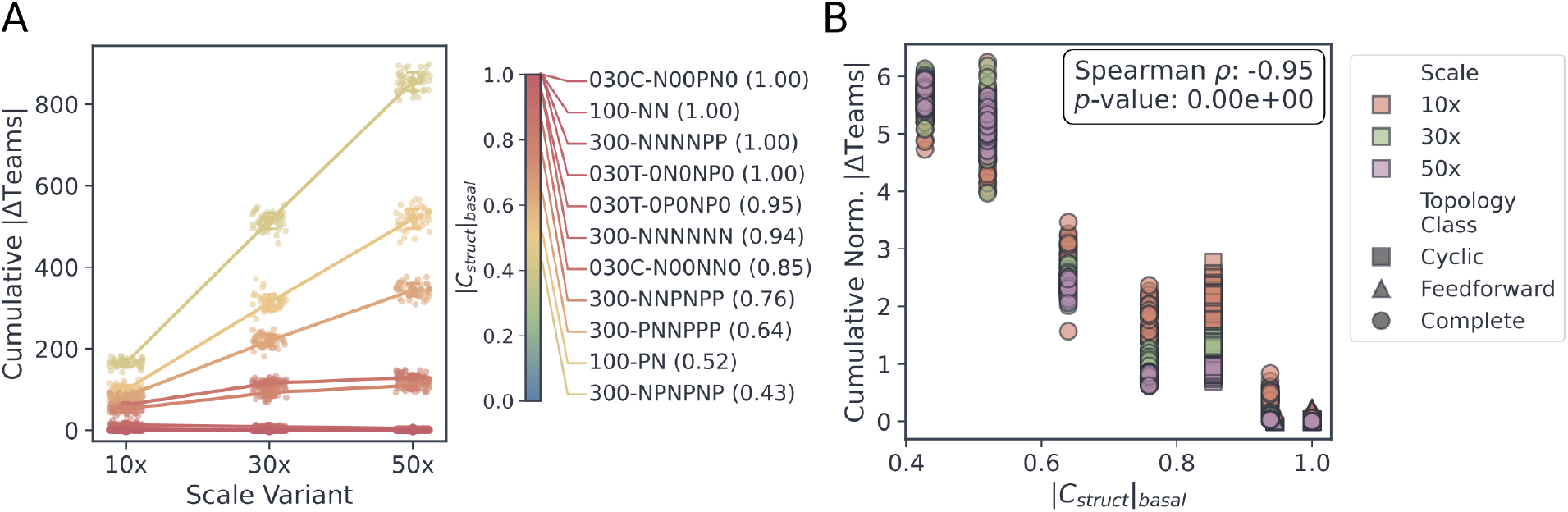
Cumulative error distributions and baseline coherence correlations for SA networks. (**A**) Comparison of the distribution of Cumulative Absolute Error (CAE) across scaling factors (10×, 30×, 50×) for all scaled SA circuits. (**B**) Scatter plot correlating Normalized Cumulative Absolute Error (Norm-CAE) against the absolute structural coherence of the circuit (|C_struct_|) for the SA networks. Individual markers represent unique circuit configurations across all scales and topology classes. The Spearman rank correlation coefficient (*ρ*) and associated p-value are indicated in the figure inset.

Figure S5A). Notably, this scale-dependent susceptibility was most pronounced in circuits with lower absolute structural coherence, which suffered the highest cumulative errors. To enable scale-agnostic comparisons, we normalized this cumulative error by the total number of nodes (*N*) to yield the Normalized Cumulative Absolute Error (Norm-CAE):

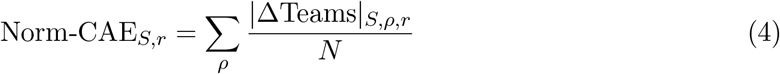

Correlating this scale-normalized error against absolute basal coherence revealed a highly predictable, statistically significant negative relationship across both SA (Spearman *ρ* = −0.95, *p <* 0.05) and NS networks (Spearman *ρ* = −0.68, *p <* 0.05) (Figure 3B, Figure S5B). These findings establish that a circuit’s initial structural coherence is a major determinant of its robustness when scaled into sparse, high-dimensional topologies.

### 2.3 Topological Architecture and Network Scale Modulate Resilience to Regulatory Disruption

Consistent with individual sparsification trajectories, complete and cyclic topologies exhibited the highest mean Norm-CAE and variance (Figure 4A), indicating high susceptibility to density-driven regulatory disruption. Conversely, feedforward architectures demonstrated strong resilience, maintaining structural integrity independent of their baseline structural coherence. These topological constraints were preserved across both the SA and NS network ensembles (Figure S6A).

**Figure 4:**
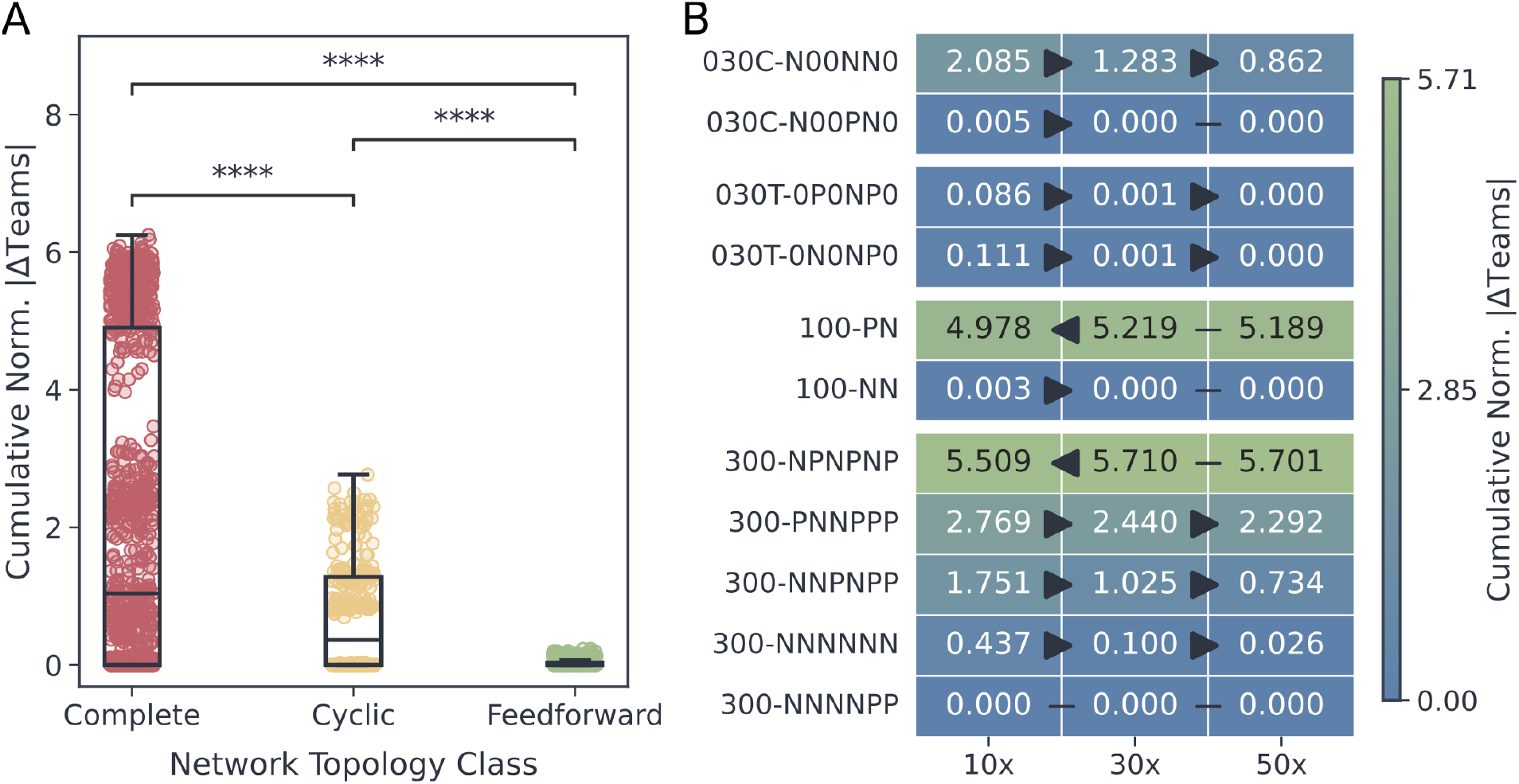
Topological resilience and scale-dependent buffering of self-activation networks under sparsification. (**A**) Box plots comparing the distribution of normalized cumulative absolute error (Norm-CAE) across topological classes for the SA networks. Statistical comparisons between distributions were performed using the Mann–Whitney U test; significance levels are denoted as ns (*p >* 0.05), * (*p* ≤ 0.05), ** (*p* ≤ 0.01), *** (*p* ≤ 0.001), and **** (*p* ≤ 0.0001). (**B**) Heatmap displaying the mean Norm-CAE values for each circuit across network scaling factors. Connecting lines between adjacent cells indicate statistical relationships; arrows heads point from a higher mean to a statistically significant lower mean, whereas flat lines without arrowheads denote no statistically significant difference according to Mann-Whitney U test.

Analyzing Norm-CAE across 10×, 30×, and 50× scaling factors revealed a scale-dependent buffering effect (Figure 4B, Figure S6B). For highly coherent circuits, error remained negligible across all scales. For circuits with intermediate baseline coherence, larger network sizes effectively buffered against coordination loss, yielding lower normalized errors. However, this trend reversed for highly incoherent base circuits (e.g., 100-PN, 300-NPNPNP). In these intrinsically incoherent architectures, particularly within the NS ensemble, higher scaling factors actively exacerbated regulatory disruption under sparsification.

Because self-activation alters baseline structural coherence, we compared Norm-CAE distributions between the SA and NS ensembles at the highly susceptible 10× scale (Figure 5A). The impact of direct auto-regulation depended strictly on the underlying circuit logic. For positive-feedback-driven circuits (e.g., 100-NN), self-activation stabilized the team structure, reducing error compared to NS variants. Conversely, in negative-feedback-dominated circuits (e.g., 100-PN), adding positive auto-regulation conflicted with the intrinsic inhibitory structure, drastically exacerbating regulatory disruption. To systematically quantify this condition-dependent susceptibility, we calculated the normalized difference of the Norm-CAE values between the NS and SA networks in mean Norm-CAE for each circuit:

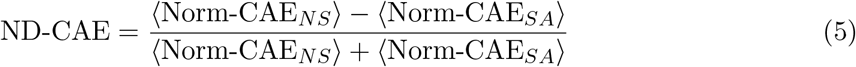

**Figure 5:**
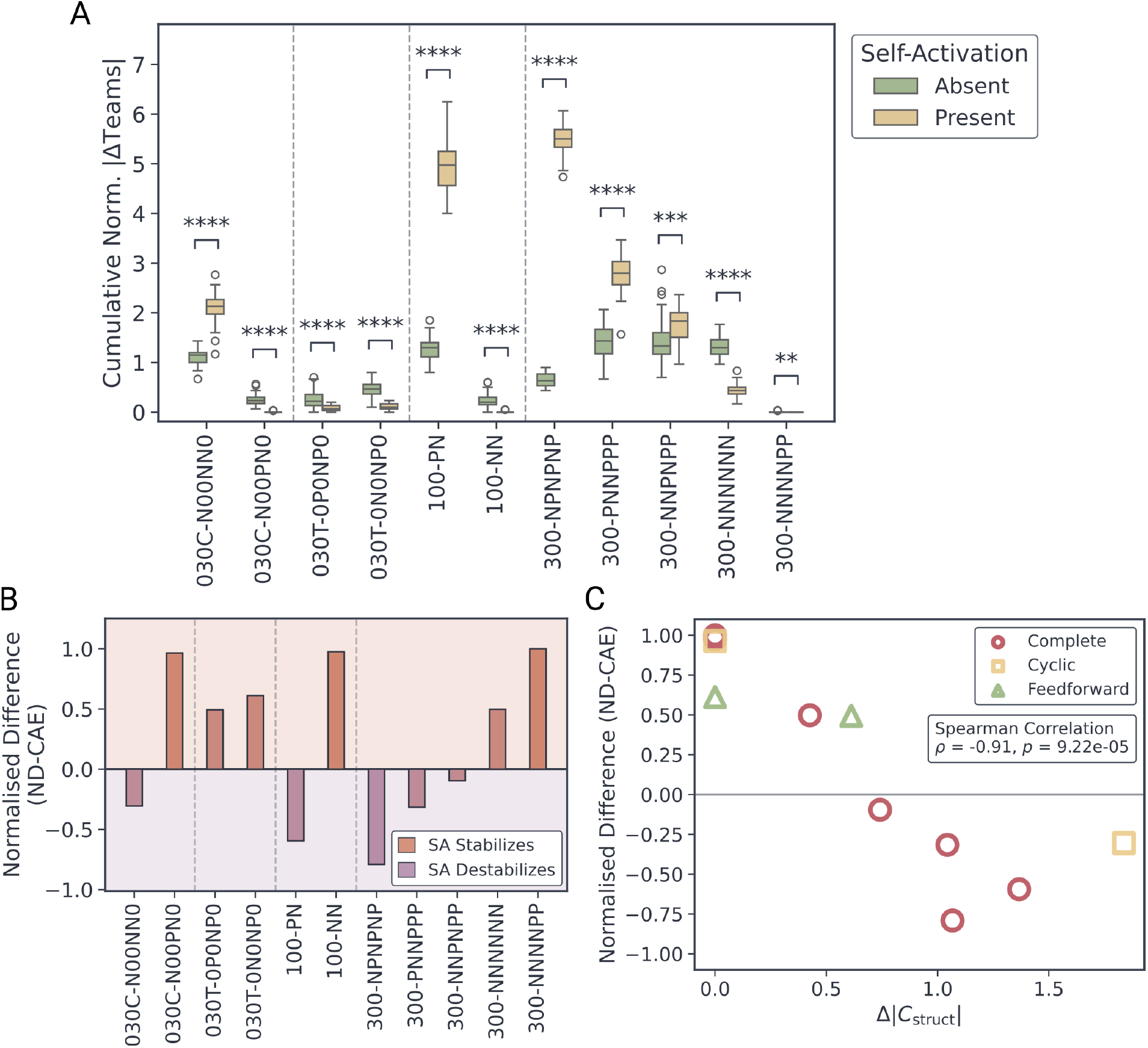
Impact of self-activation on the robustness of scaled circuits under sparsification. (**A**) Comparison of Norm-CAE distributions between NS and SA networks for individual base circuits at the 10× scale. Circuits are grouped by topological classes, separated by vertical dotted lines. Statistical comparisons between distributions were performed using the Mann–Whitney U test; significance levels are denoted as ns (*p >* 0.05), * (*p* ≤ 0.05), ** (*p* ≤ 0.01), *** (*p* ≤ 0.001), and **** (*p* ≤ 0.0001). (**B**) Bar plot of the normalized difference in mean Norm-CAE (ND-CAE) between NS and SA networks. Shaded background regions denote whether self-activation stabilizes (ND-CAE *<* 0) or destabilizes (ND-CAE *>* 0) the network topology. (**C**) Scatter plot correlating ND against the absolute structural coherence shift (Δ|C_struct_|) induced by self-activation. Individual markers differentiate topological classes. The Spearman rank correlation coefficient (*ρ*) and p-value are shown in the inset legend.

A positive ND-CAE indicates greater susceptibility in the NS variant, whereas a negative ND-CAE highlights greater vulnerability in the SA variant. We evaluated ND-CAE against the self-activation-induced shift in absolute structural coherence (Δ|*C*_struct_| = |*C*_struct, SA_| − |*C*_struct, NS_|). This revealed a strong negative correlation (Spearman *ρ* = −0.91, *p <* 0.05) (Figure 5B-C). Ultimately, enforcing localized positive auto-regulation stabilizes architectures governed by positive feedback, but actively disrupts team structures in circuits dominated by negative feedback loops.

### 2.4 GRNs Maximize Regulatory Flexibility Through Team Organization

To evaluate how regulatory alignment manifests in biological contexts, we applied the coherence framework to six whole-organism GRNs from the Abasy Atlas [42]. We utilized meta-curated, signed, and directed networks strictly filtered for high-confidence evidence to capture genome-wide regulatory behavior, avoiding the localized biases inherent in process-specific sub-networks used to model biological phenomenon that offer an incomplete representation of system-wide coordination. This generated a dataset of GRNs spanning diverse levels of coverage and completeness (Table 1). Across all organisms, the distribution of the matrix elements *C*_*ij*_ within the computed coherence matrices was predominantly bimodal, reflecting a baseline of highly coherent connectivity (Figure 6A, Figure S7). To investigate the topological constraints shaping this distribution, we compared wild-type (WT) GRNs against randomized null models. In these randomized networks, the incoming and outgoing node degree distributions as well as the global regulatory-sign distribution were strictly preserved, while the specific source-to-target wiring was shuffled (see Supplementary Methods). The randomized topologies consistently generated sharper, higher density peaks at the absolute coherence extremes of −1 and +1 compared to the WT networks (Figure 6A). Upon identifying teams within these coherence matrices we observed that the WT architectures partitioned into a significantly greater number of teams than their corresponding randomized counterparts (Figure 6B). This elevated team count demonstrates that whole-organism regulatory networks favor regulatory modularity over global co-expression. The higher frequency of mixed, lower-coherence interactions observed in WT architectures might provide increased regulatory degrees of freedom, allowing targeted gene sets to respond independently to diverse signals while maintaining the baseline local coordination required to act in concert.

**Table 1.** Prokaryotic gene regulatory networks used for analysis.

| **Organism** | **Coverage** | **Completeness** |
|---|---|---|
| *Corynebacterium glutamicum* | 71.7% | 42.6% |
| *Mycobacterium tuberculosis* | 62.1% | 67.3% |
| *Bacillus subtilis* | 58.2% | 49.4% |
| *Escherichia coli* | 51.3% | 56.0% |
| *Pseudomonas aeruginosa* | 18.4% | 13.7% |
| *Streptomyces coelicolor* | 5.2% | 2.3% |

**Figure 6:**
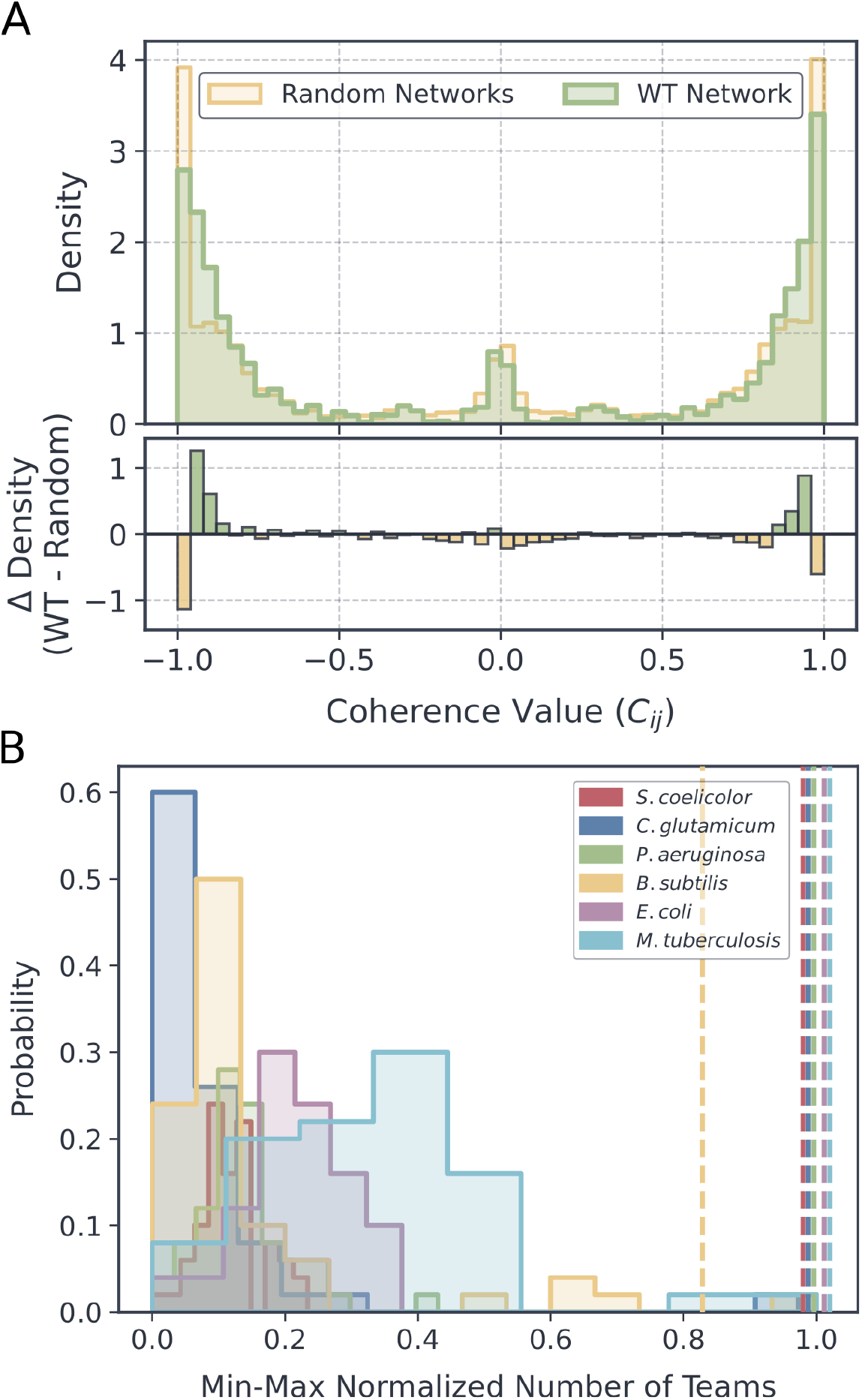
Coherence distributions and team organization structure in whole-organism GRNs. (**A**) Probability density distribution of coherence values comparing aggregated wild-type (WT) whole-organism GRN of *E*.*coli* against degree-preserving randomized networks. The bottom subpanel depicts the density difference (Δ Density = Shuffled - WT). (**B**) Box plots comparing, for each organism, the min-max normalized total number of identified teams in WT networks (horizontal dashed lines) against the distribution of number of teams observed in their randomized counterparts. For *B. subtilis*, only 1 out of 50 random networks exhibited fewer teams than observed in the WT networks. For the other four bacterial GRNs, all 50 randomized networks exhibited less or equal number of teams than the respective WT network.

### 2.5 Functional Compartmentalization and the Centralization of Higher-Order Interactions in GRNs

To characterize the functional roles of genes within these coordinated teams, we evaluated the network architectures using established hierarchical layering frameworks [12, 13, 14]. This structural decomposition served two primary objectives: delineating how structural coherence is distributed across biologically annotated functional modules, regulatory teams, and interaction circuits, and extracting the empirical block-densities required to parameterize our subsequent synthetic hierarchical networks. By utilizing the coherence matrix to explicitly distinguish absolute topological disconnection (represented by undefined values) from active regulatory coherence, we classified structurally integrated genes according to their directional connectivity. Nodes exhibiting exclusively outgoing coherence were designated as input genes, those with exclusively incoming coherence as output genes, and those possessing both as middle-layer genes (see Supplementary Methods).

Consistent with previously reported results [12, 13, 14], this classification revealed a highly skewed, globally feed-forward architecture: input (4.7% ± 4.1%) and middle (8.5% ± 3.2%) genes were scarce, while terminal output genes (86.8% ± 7.0%) dominated the topology (Figure S8A). Inter-layer density quantification confirmed that the middle layer maintains the highest internal connectivity (Figure S8B; Figure S9). This structural template directly contextualizes the sparsification dynamics observed in our synthetic Erdős–Rényi (ER) ensembles. Because local feed-forward topologies natively demonstrated superior structural resilience against density-driven regulatory disruption, establishing a globally feed-forward macroscopic hierarchy provides a scalable topological mechanism to preserve systemic coordination under extreme biological sparsity (Figure 4A, S6A). However, this macroscopic perspective does not address the localization of well-documented regulatory decision-making motifs [15, 8, 16, 17, 18, 19, 20], many of which fundamentally rely on mutual or cyclic feedback, nor does it resolve how these discrete local structures integrate into broader functional modules and teams of activating nodes.

Further analysis of functional modules defined by the Abasy Atlas [42] revealed that, after excluding global regulatory machinery (*>* 95th percentile size), module sizes followed a long-tailed distribution characterized by a median of fewer than ten genes (Figure S10A) (see Supplementary Methods). While the global network architecture exhibits a fewer numbers of input and middle nodes, the relative composition within individual functional modules demonstrated greater variability, though it still retained a pronounced skew toward output genes (Figure S10B). Across all networks, intra-module connectivity consistently exceeded inter-module coupling, confirming that regulatory interactions preferentially cluster within distinct biological processes (Figure S11A, B). Despite this dense inter-module interaction, systemic inter-module coordination was maintained by mostly relying on directed interactions routed through the restricted middle layer (Figure S11A-ii).

To characterize the functional stratification imposed by these topological constraints, we performed layer-specific Gene Ontology (GO) enrichment analysis (see Supplementary Methods). The sparse input and middle layers exhibited significantly higher global odds ratios than the output layer (Figure S12A, B), indicating tight functional specialization in the upper strata contrasting with the pronounced heterogeneity of the output layer. Furthermore, individual functional modules were not restricted to specific layers, confirming that discrete biological processes span the entire regulatory hierarchy (Figure S12C). To characterize specific layer functions, significant GO terms were ranked via a specificity-weighted Combined Score (see Supplementary Methods). This analysis confirmed a strict functional stratification: the input layer acts as an environmental interface enriched for signal transduction and minor TF activity (Figure S13); the compact middle layer centralizes integration via an overwhelming enrichment of TF activity and DNA-binding functions, a functional shift further corroborated by a substantially higher odds ratio of TFs localizing specifically to this layer (Figure S12D); and the large output layer remains depleted of regulatory annotations, functioning as a heterogeneous effector compartment composed of metabolic and enzymatic machinery that executes middle-layer driven decisions (Figure S13).

To evaluate localized regulatory logic, we next analyzed the enrichment of 2- and 3-node directed, signed circuits, categorized as feedforward, cyclic, complete, or complex (composite circuits that combine mutual, cyclic and feed-forward interactions) (see Methods) (Figure S1). Feedforward circuits (030T), complete mutual dyads (100) and circuits containing mutual dyads (e.g., 111U, 120D) were significantly enriched, whereas unidirectional dyads (010), cyclic circuits (021C, 030C), and circuits with two pairs of mutual interactions (201, 210) were depleted (Figure 7A). Despite the globally feed-forward architecture, the enrichment of mutually connected motifs highlights dense local connectivity within the core. Notably, the fully connected triad (300) was absent; the only enriched complete topology was the mutual dyad (100). Enriched circuits generally exhibited high absolute structural coherence, irrespective of topology class, but, no clear trend was present between a circuit’s enrichment and its structural coherence (Figure S14A, B). Circuits containing recursive feedback (complete, cyclic, complex) consistently harbored multiple TFs (≥ 2), whereas feedforward circuits typically contained one (Figure 7B, Figure S15). Topologically restricted to the middle layer by definition, these multi-TF, feedback-heavy motifs are physically sequestered within the dense regulatory core, where they span diverse functional modules and teams to integrate systemic signals (Figure 7B).

**Figure 7:**
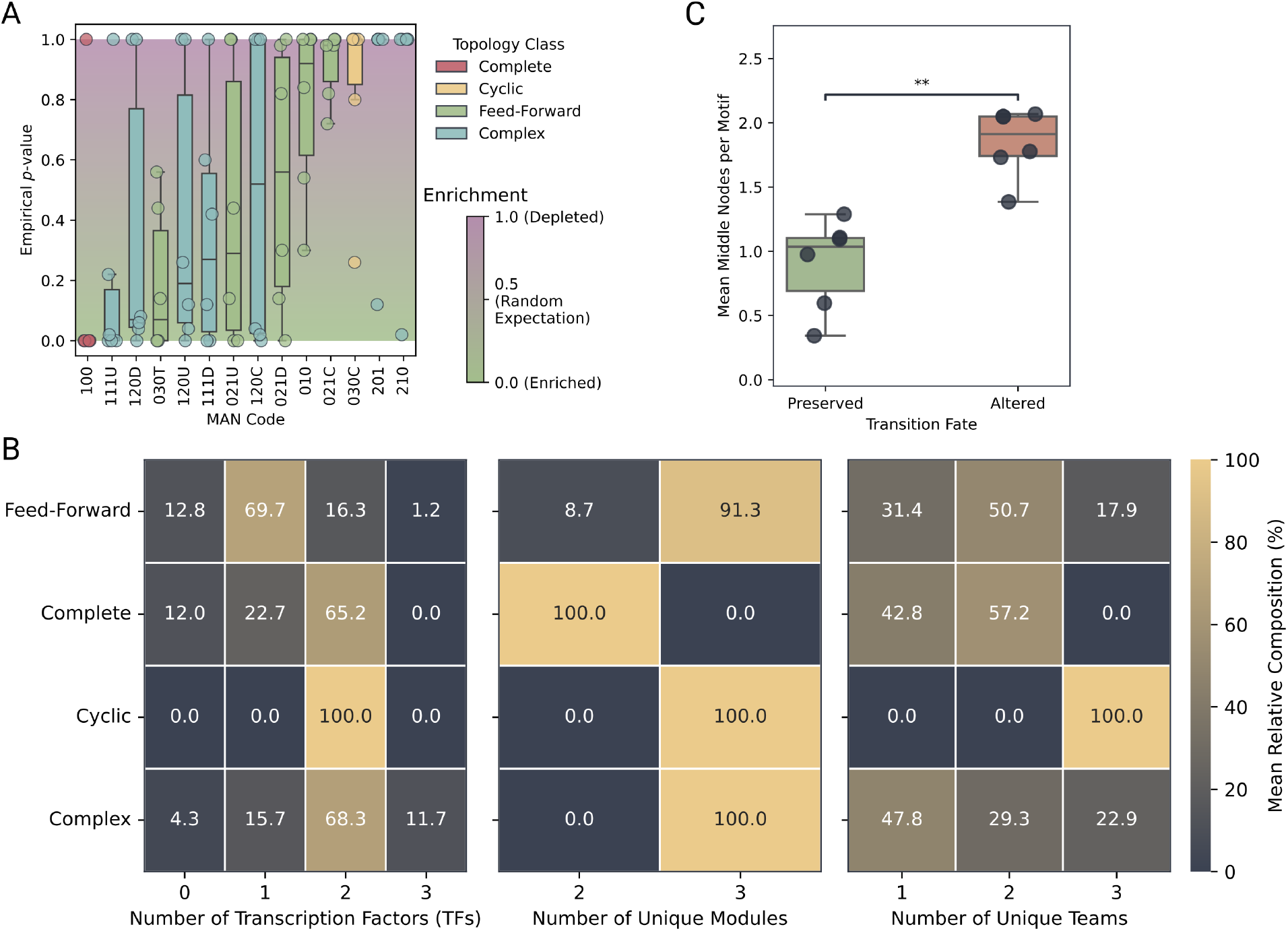
Motif enrichment, structural composition, and topological stability. (**A**) Box plots displaying the distribution of empirical p-values for the MAN codes observed in the whole-organism networks. (**B**) Heatmaps detailing the structural composition across the four topological classes of circuits. From left to right, the panels display the relative distribution of the number of transcription factors, unique functional modules, and unique teams present within each circuit class. Values denote the mean percentage of circuits exhibiting the specified count, with each row summing to 100% within individual heatmaps. (**C**) Box plots comparing the mean number of middle nodes within circuits that remain structurally preserved versus those that are altered when transitioning from the adjacency matrix to the coherence matrix. Statistical comparisons between distributions were performed using the Mann–Whitney U test; significance levels are denoted as ns (*p >* 0.05), * (*p* ≤ 0.05), ** (*p* ≤ 0.01), *** (*p* ≤ 0.001), and **** (*p* ≤ 0.0001).

Evaluating topological stability when transitioning from the discrete adjacency to the continuous coherence matrix, which integrates higher-order indirect paths, revealed that feedforward and complete topologies largely preserved their structures (Figure S16A). Furthermore, transcription factors were more frequently involved in these altered motifs than in preserved ones, highlighting their greater participation in feedback loops (Figure 7C). These findings indicate that these topological structures operate not as isolated local circuits, but as dynamic, higher-order integrative hubs. They frequently embed within broader decision-making architectures responsible for multi-signal integration (e.g., 111U) and coordinated signal dissemination (e.g., 120D, 120U) [19, 43]. By restricting these small but dense feedback-accumulating topologies to the middle stratum, the hierarchy enforces a centralized routing architecture. This systematic aggregation of upstream inputs, integrated via dense regulatory crosstalk within the middle layer, generates broadly distributed and coherent instructional signals. Such centralized coordination allows the network to drastically scale its terminal effector layer under stringent global sparsity, irrespective of baseline regulatory coherence.

### 2.6 Hierarchical Architecture Limits Density-Driven Team Disruption

To explicitly test whether a restricted, densely connected middle layer preserves structural coordination, we constructed synthetic hierarchical (HI) networks mimicking biological proportions: 5% input, 5% middle, and 90% output nodes. Feedforward base circuits were excluded, as their topology is inherently hierarchical. Unlike the ER models, sparsification was evaluated by modulating only the input-to-output connection density (10% to 100%). This targeted modulation isolates inter-layer signal integration and satisfies structural necessity; a 5% middle layer contains too few nodes to be globally sparsified without severing regulatory links entirely. Due to these strict block constraints, a 100% input-to-output density in an HI network corresponds to a maximum global density of only 8.7% (see Methods and Supplementary Methods).

Evaluating the scale-normalized Cumulative Absolute Error (Norm-CAE) for the HI ensembles revealed that, similar to ER networks, lower absolute structural coherence generally corresponded to higher cumulative regulatory disruption (Figure 8A, Figure S17A). However, correlating Norm-CAE against basal |C_struct_| yielded significantly attenuated correlations compared to ER networks: −0.51 (*p <* 0.05) for the SA ensemble and −0.31 (*p <* 0.05) for the NS ensemble (Figure S17B, i-ii). This attenuation is consistent with hierarchical networks being less susceptible to the structural coherence of their base circuits, effectively buffering against the regulatory disalignment that drives ER network failure.

**Figure 8:**
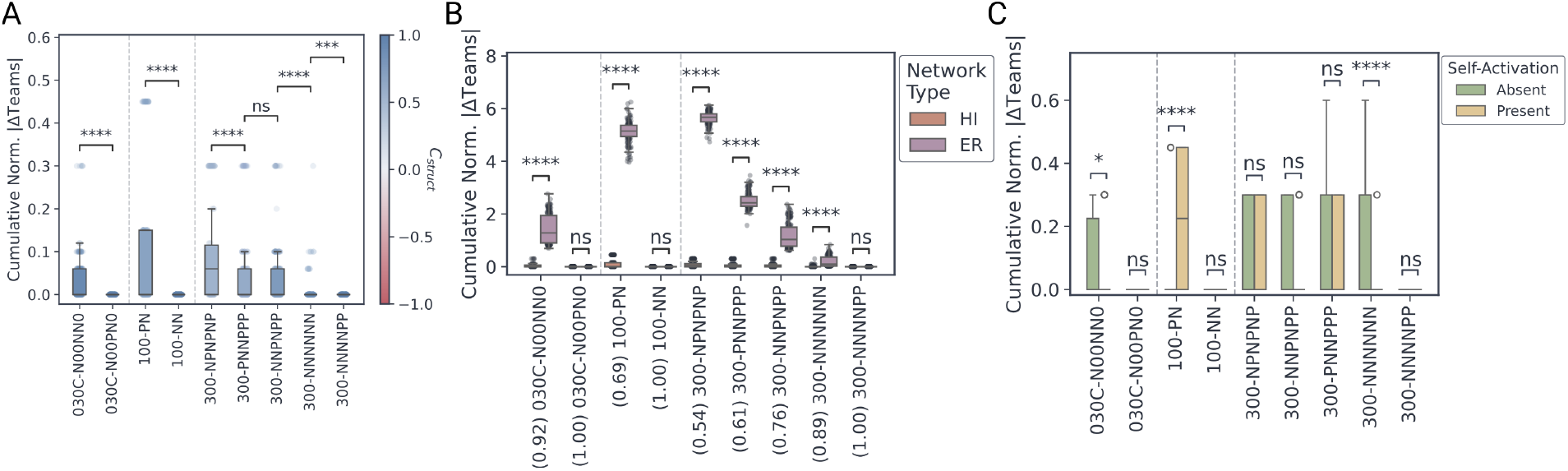
Structural robustness of hierarchical networks under sparsification. (**A**) Box plots of the distribution of Norm-CAE values across all scaling factors for individual base circuits of the SA hierarchical networks. Circuits are grouped by topological classes, separated by vertical dotted lines. Box colors map to the absolute structural coherence (|C_struct_|) of the corresponding base circuit. (**B**) Box plots comparing the Norm-CAE distributions between ER and HI SA networks for individual base circuits. X-axis labels denote the specific circuit MAN code and its baseline C_struct_ in parentheses. (**C**) Box plots comparing Norm-CAE distributions between NS and SA hierarchical networks for individual base circuits at the 10× scale. Circuits are grouped by topological classes, separated by vertical dotted lines. Statistical comparisons between distributions were performed using the Mann–Whitney U test; significance levels are denoted as ns (*p >* 0.05), * (*p* ≤ 0.05), ** (*p* ≤ 0.01), *** (*p* ≤ 0.001), and **** (*p* ≤ 0.0001).

Direct comparisons of Norm-CAE distributions confirmed this buffering effect. Across all scales, HI networks exhibited markedly lower cumulative absolute errors than their ER counterparts (Figure 8B). Crucially, the stabilization magnitude scaled inversely with absolute structural coherence: circuits with the lowest coherence, which suffered higher regulatory disruption in ER contexts, remained highly constrained within HI topologies. This differential stabilization, observed consistently in both SA and NS networks, demonstrates that HI architectures maintain cohesive team structures across diverse regulatory constraints despite operating at extreme global sparsity.

Finally, comparing the 10× HI SA and HI NS ensembles revealed that self-activation continues to exert a context-dependent effect on structural stability (Figure 8C). As in ER models, forced SA disrupted negative-feedback-dominated circuits (e.g., 100-PN) while stabilizing positive-feedback architectures. Because hierarchical topology buffers overall baseline regulatory disruption to near-zero, global correlation between structural coherence shifts (Δ|C_struct_|) and normalized difference lacked statistical significance (Figure S17D).

Collectively, these findings elucidate the structural strategy GRNs may employ to achieve scalable coordination. By embedding intrinsically conflicting or low-coherence regulatory logic strictly within a dense middle layer, and confining global signal propagation to a feedforward hierarchical architecture, biological systems structurally buffer against coordination loss. This allows them to maintain stable regulatory teams independent of intrinsic circuit coherence, resolving the dual constraints of system scale and network sparsity.

### 2.7 Validation of Coherence-Based Modules Using Transcriptomic iModulons

While our analyses of synthetic architectures demonstrate that hierarchical buffering preserves structural coordination against extreme sparsity, these findings rely on static network topologies and their derived behavior. To determine whether the coherence formalism captures biologically active regulatory relationships, we evaluated whether topology-derived coherence values encode predictive information about empirical gene expression states.

We compared coherence-based predictions against functional gene modules defined by *Es-cherichia coli* iModulons [27]. The iModulon framework decomposes expression profiles into independent components, establishing a data-driven reference for coordinated gene responses.

We first performed a coarse-grained comparison, examining the overlap between iModulon member genes and those connected by nonzero coherence values (see Methods). Although a positive association existed between iModulon size and the number of retained genes, coherence consistently underestimated module sizes (Figure 9A). This under-reporting likely reflects inherent incompleteness in the static regulatory network and the inclusion of weakly weighted, lower-threshold genes within the empirical iModulon definitions.

**Figure 9:**
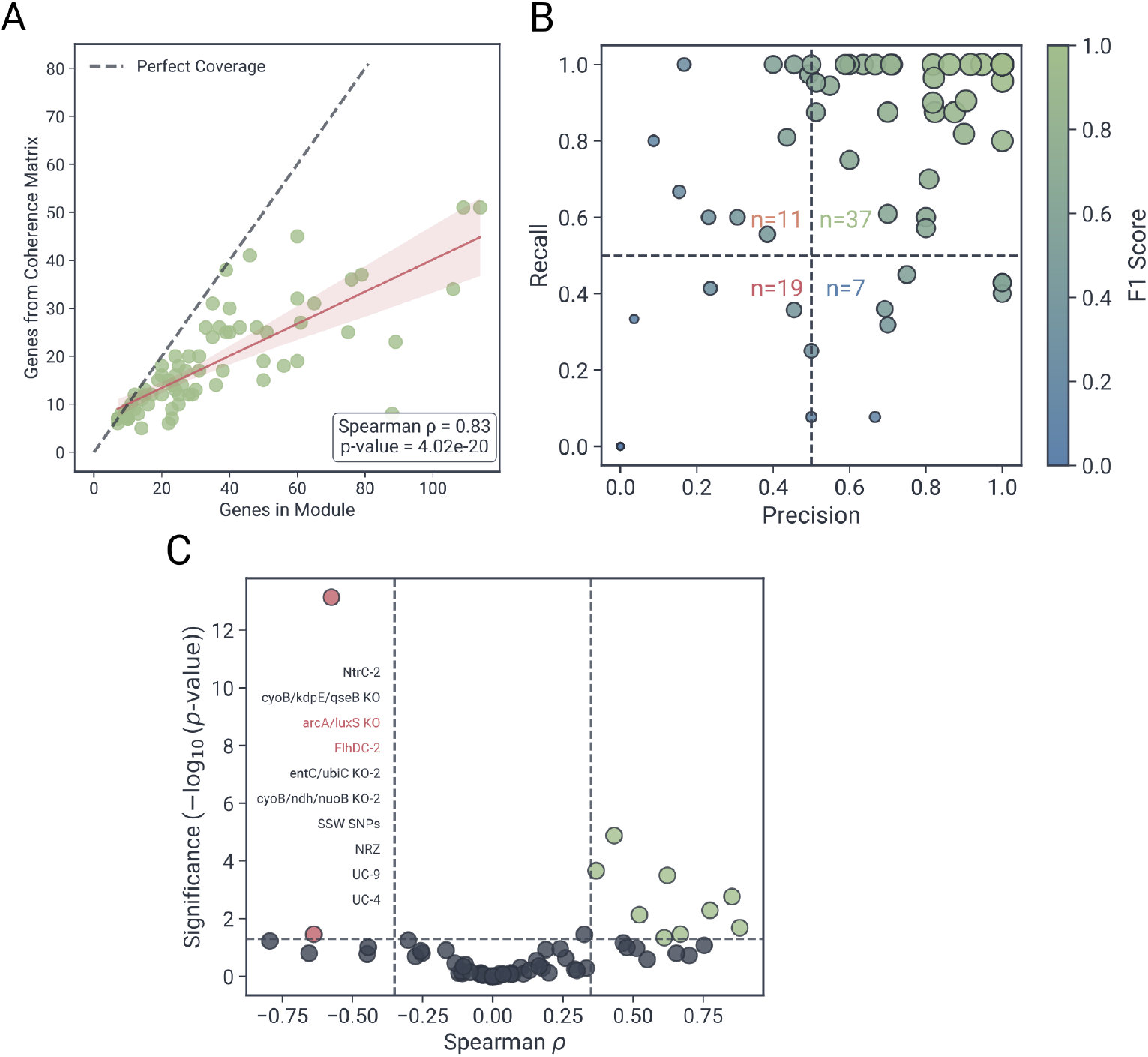
Validation of coherence-based predictions against transcriptomic modules. (**A**) Scatter plot detailing module coverage, comparing the total number of genes in an identified iModulon against the number of those genes present within the coherence matrix. The dashed line denotes perfect coverage; the solid line represents the linear regression fit with the confidence interval shaded. (**B**) Scatter plot evaluating classification performance for predicting the sign of gene-gene interactions within each iModulon, mapping recall against precision. Individual markers represent specific modules, color-coded by their F1 score. Dashed lines indicate the 0.5 performance threshold for both metrics. (**C**) Volcano plot of the Spearman rank correlation between the magnitude of coherence values and iModulon interaction weights, against statistical significance. Vertical dashed lines denote correlation thresholds of ±0.3; the horizontal dashed line marks the significance threshold (*p* = 0.05). Points are color-coded by significance and correlation direction: significantly positive (green), significantly negative (red), and non-significant (grey). Red text labels identify the significant, negatively correlated iModulons.

We next evaluated the ability of the coherence matrix to predict the sign of gene–gene interactions. For all pairwise combinations within an iModulon, we computed the product of their component weights to determine whether the genes contribute in a concordant or antagonistic manner. Comparing these empirical interaction signs against the signs of the corresponding coherence values (see Methods) yielded high precision and recall across most modules, reflected in elevated F1 scores (Figure 9B). These results demonstrate that coherence predicts, to a large extent, the directionality of coordinated gene expression solely on the basis of the static regulatory topology.

We further investigated whether coherence captures the relative magnitudes of these interactions by correlating the pairwise products of iModulon weights with the corresponding coherence values. While global correlations across the full dataset were weak, a subset of modules exhibited statistically significant positive correlations, aligning topological coherence strength with graded expression contributions (Figure 9C). This alignment is notable given that coherence values are strictly bounded (±1), whereas iModulon weights occupy a continuous range. However, modules exhibiting poor coherence predictions were predominantly associated with global stress responses, gene knockouts, or highly condition-specific programs. In such contexts, the active regulatory architecture is dynamic. Because coherence is derived from a static structural scaffold, it cannot account for condition-dependent inactivation or rewiring of network components. To test this limitation explicitly, we analyzed gene expression variance under varying global stress conditions [44] and found no significant correlation between input coherence and absolute log_2_ fold changes (Figure S18; see Methods). This lack of association highlights that static coherence topologies do not capture the dynamic variation driven by global regulatory rewiring.

## 3 Discussion

Understanding how gene regulatory networks (GRNs) maintain systemic coordination despite extreme sparsity and combinatorial complexity remains a central challenge. While classical graph-theoretic analyses identify key organizational features like degree distributions [5, 6, 7], modularity [8, 9, 10, 11], and hierarchical levels [12, 13, 14], their reliance on unsigned, localized connectivity obscures how diverse regulatory signals propagate. To address this, we introduce the coherence matrix. By integrating indirect paths, this approach moves beyond traditional sign-agnostic metrics to explicitly quantify the consistency and sign of regulatory influence. Building on structural balance frameworks [23, 45, 25, 24, 46] and influence matrices [21, 22], the coherence matrix yields strictly bounded, interpretable measures of regulatory balance. Crucially, by propagating undefined values (NaNs) for unlinked pairs, it prevents conflating topological disconnection with balanced regulatory conflict. This dual advantage resolves the ambiguity of unbounded metrics, enabling direct comparisons across highly sparse networks.

Analyzing synthetic Erdős–Rényi (ER) networks demonstrated that scaling low-coherence regulatory circuits requires significantly higher connection densities to maintain cohesive team structures, whereas highly coherent architectures naturally resist sparsification. This baseline resilience is dictated by topological class: feed-forward architectures resist density-driven disruption, while cyclic and complete architectures containing feedback loops are highly susceptible. Network scale modulates these penalties—larger scaling factors protect moderately coherent circuits, while extreme scaling exacerbates disruption in intrinsically conflicting architectures. Furthermore, localized self-activation uniquely stabilizes positive-feedback-driven networks while actively disrupting regulatory alignment in those dominated by negative feedback.

Applying this framework to six whole-organism GRNs revealed how whole-organsim systems deploy these constraints within a conserved three-strata hierarchy: a sparse input layer, a dense intermediate core of transcriptional regulators, and a broad effector output layer [12, 13, 14]. Coherence analysis demonstrated that biological GRNs partition into significantly more teams than randomized null models. Rather than operating as rigid monoliths, whole-organsim architectures favor structural modularity, maximizing regulatory degrees of freedom to process diverse signals independently. Furthermore, tracking regulatory integration across varying paths revealed that complex, low-coherence motifs persist despite their scaling penalties strictly because they are confined to the small, dense intermediate core. This core centralizes regulatory crosstalk, routing systemic coordination across independent modules while preserving global sparsity. By confining strongly connected components, recursive feedback loops, and the structural elements of the feedback vertex set (FVS) [47, 48, 49, 50] to this central regulatory hub, biological networks scale coherent and incoherent regulatory interactions, maintaining dense connectivity exclusively where signal integration is required.

Synthetic hierarchical (HI) networks mimicking these scale-invariant proportions confirmed the functional consequences of this architecture. Unlike ER baselines, HI networks constrained regulatory disruption (Norm-CAE), maintained cohesive team structures across all parameters, and neutralized the destabilizing effects of local auto-regulation—all while operating at extreme global sparsity (maximum effective density of 8.7%). This demonstrates that macroscopic hierarchical topology explicitly neutralizes density-driven scaling penalties. By embedding conflicting regulatory logic within a dense core and restricting global propagation to a feed-forward scaffold, biological systems decouple team cohesion from baseline structural coherence, simultaneously maximizing scale and minimizing network sparsity.

Transcriptomic validation against *E. coli* iModulons [27] corroborated the biological relevance of this framework. Computed exclusively from static topology, the coherence matrix successfully predicted the direction of co-regulation (concordant versus discordant) within functional modules, and occasionally aligned with relative expression magnitudes. Deviations from topological predictions emerged primarily during global stress or gene knockouts, where transcriptional rewiring dynamically shifts the active regulatory architecture. These context-dependent shifts motivate extensions for the coherence framework: integrating continuous transcript abundance, chromatin accessibility, or Boolean logic constraints [51] as edge weights to transition from static structural baselines to dynamic descriptors of context-aware regulatory potential.

Collectively, these results establish structural coherence as a complementary framework for analyzing signed GRNs. By moving beyond traditional sign-agnostic metrics to explicitly quantify the direction and sign of regulatory influence, this approach links localized regulatory logic to the emergence of system-wide coordination under sparsity, offering a robust tool for decoding complex biological networks.

## 4 Methods

### 4.1 The Coherence Matrix

We considered a signed, directed gene regulatory network represented by an adjacency matrix

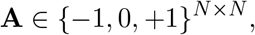

where *A*_*ij*_ = +1 denotes activation of node *i* by node *j, A*_*ij*_ = −1 denotes inhibition, and *A*_*ij*_ = 0 indicates the absence of a direct regulatory interaction.

To quantify the regulatory influence of node *j* on node *i* through paths of a fixed length *l*, we examined the *l*-th power of the adjacency matrix, **A**^*l*^. The corresponding matrix elements

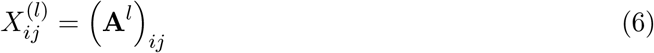

represent the net signed influence accumulated over all walks of length *l* from *j* to *i*. Specifically,

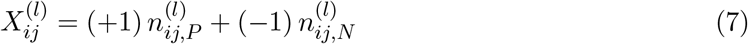

where 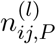 and 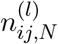 denote, respectively, the number of positive (activating) and negative (inhibitory) walks of length *l* from node *j* to node *i*. A walk is considered positive when it includes an even number of inhibitory edges, and negative when the number of inhibitory edges is odd.

We also quantified the total number of walks independently of regulatory sign by considering powers of the absolute adjacency matrix |**A**|, defined elementwise as |**A**|_*ij*_ = |*A*_*ij*_|. The matrix power

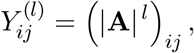

counts the total number of walks of length *l* from node *j* to node *i*, irrespective of the signs of the edges along the walk. So for each element *Y*_*ij*_,

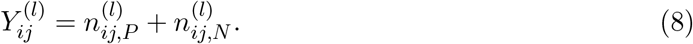

For node pairs connected by at least one walk of length 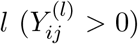, we defined a length-specific coherence matrix 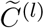 with elements,

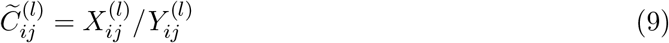

This maps purely activating interactions to +1, purely inhibitory interactions to −1, and an equal balance to 0. If no walk of length *l* exists between a node pair 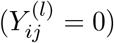, the corresponding entry 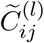 is left undefined (NaN).

To integrate regulatory influence across multiple topological scales, we aggregated the centered coherence matrices over all path lengths up to a maximum *L*, weighting contributions from longer paths by a factorial decay that suppresses increasingly indirect effects. The final Coherence Matrix *C* is defined element-wise as

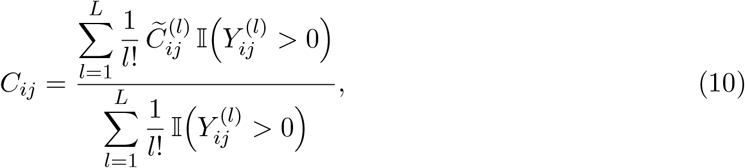

where I(·) is an indicator function that excludes undefined (NaN) entries arising from path lengths for which no walks exist between the node pair. This represents a conditional normalization where only path lengths for which at least one walk is present 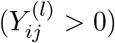 contribute to both the numerator and denominator. As a result, *C*_*ij*_ is a weighted average over the subset of path lengths that actually connect nodes *j* and *i*. For this study, we evaluated paths up to a maximum length of *L* = 10. Because the factorial weighting term (1*/l*!) decays rapidly (e.g., 1*/*10! ≈ 2.7 × 10^−7^), the relative contribution of longer paths becomes vanishingly small. Consequently, *L* = 10 serves as a highly conservative upper bound that fully captures long-range indirect interactions while preventing numerical instabilities arising from computing high matrix powers in large networks. Depending on the required precision, even shorter maximum walk lengths can be utilized without significantly altering the computed coherence values.

The sign of each element in *C, C*_*ij*_, reflects whether activating or inhibitory walks dominate the influence of node *j* on node *i*, whereas the magnitude |*C*_*ij*_| measures the degree of regulatory coherence across all paths linking the two nodes.

#### 4.1.1 Team identification via Coherence Matrix partitioning

We identified functional “teams” as maximal subsets of genes exhibiting mutual regulatory activation using a multi-stage clustering and refinement algorithm applied to a discretized signed coherence matrix *S*, where *S*_*ij*_ = sign(*C*_*ij*_) ∈ {−1, 0, 1}. The algorithm isolates nodes lacking self-activation, initializes core groups based on strict regulatory compatibility of no inhibitions within a team, and refines them through a sequential greedy optimization that maximizes positive internal connectivity. A detailed formalization of the algorithm is provided in the supplementary section A.1. The final requirement for the nodes in the identified teams to be weakly connected was enforced only for Erdős–Rényi networks, as applying it to the sparse, heavily skewed architectures of hierarchical and empirical networks artificially inflates team counts.

#### 4.1.2 Structural Coherence: Network-Level Coherence Measure

Beyond identifying teams, we also quantified the network’s overall regulatory consistency using a scalar structural coherence measure derived from the team-partitioned coherence matrix. Since teams are defined as subsets of nodes exhibiting mutually positive coherence, structural coherence is computed exclusively from intra-team interactions.

Let *T* = {*T*_1_, *T*_2_, …, *T*_*K*_} denote the set of teams identified for a given network. For each Team *T*_*k*_, we considered the corresponding sub-matrix of the Coherence Matrix **C** restricted to node pairs (*i, j*) such that *i, j* ∈ *T*_*k*_. Undefined entries (NaNs), corresponding to node pairs lacking any direct or indirect connectivity, are excluded from all averages.

The structural coherence *C*_struct_ was then defined as the mean of all numeric coherence values within these intra-Team blocks,

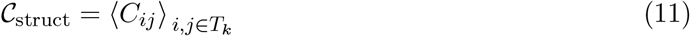

This definition yields a bounded scalar *C*_struct_ ∈ [−1, 1] that captures the average regulatory agreement within all the identified teams. *C*_struct_ is positive when activating regulation dominates within teams, near zero under mixed regulation, and negative in networks whose Team structures are governed primarily by inhibitory or negative feedback interactions.

### 4.2 Generation of Synthetic Gene Regulatory Networks

To systematically evaluate how topology and sparsity affect structural integrity, we generated synthetic GRNs by scaling a representative subset of 2-node and 3-node directed, signed base circuits. These circuits were selected to span the structural coherence spectrum and were categorized using MAN (Mutual, Asymmetric, Null) codes into fundamental topological classes: feed-forward (030T), complete (100 and 300), and cyclic (030C) [41] (Figure S1). These circuits were then expanded by scaling factors of *S* ∈ {10, 30, 50} via block expansion, effectively replacing individual nodes with coordinated groups of genes. To distinguish coordination arising from localized autoregulation versus network-wide feedback, we generated parallel ensembles with (SA) and without (NS) explicit self-activation loops in the initial circuits. For both ensembles, Erdős–Rényi (ER) type networks were generated across global edge densities from 10% to 100% using an additive edge-growth algorithm. This approach guaranteed that while supplemental edges varied independently to meet density and hierarchical constraints, the base wiring remained strictly conserved across all realizations of the same replicate and network size.

To evaluate how biological architectures modulate these scaling properties, we constructed artificial hierarchical (HI) networks using the same block-expansion framework but imposed strict scale-invariant layer constraints mirroring empirical GRNs (5% Input, 5% Middle, 90% Output). Feed-forward base circuits were excluded from this ensemble to avoid topological redundancy. Connectivity was restricted to a predominantly feed-forward architecture with fixed intermediate block densities derived from empirical constraints, while direct Input-to-Output connectivity was swept from 10% to 100%. Due to these strict architectural blockades, a maximally connected HI network under these constraints corresponds to an extreme global edge density of approximately 8.7%. Complete mathematical details regarding circuit enumeration, targeted sampling, the additive scaling procedure, and hierarchical block masking are provided in the Appendix Methods.

### 4.3 Selection and pre-processing of whole-organism GRNs

Whole-organism gene regulatory networks (GRNs) were sourced from the Abasy Atlas database [42, 52, 53, 54]. Because the coherence formalism requires signed interactions, we retained only edges with defined regulatory signs (+ or −). To ensure sufficient valid connectivity, we required that at least 20% of a GRN’s originally reported edges possess a defined sign; networks failing this threshold were excluded. To minimize spurious interactions, our analysis was strictly confined to edges categorized by the database as possessing “Strong” experimental evidence. This filtering yielded six whole-organism GRNs (1). For each GRN, we constructed a signed, directed adjacency matrix **A**, where *A*_*ij*_ ∈ {−1, 0, +1} denotes inhibitory, absent, or activating regulation, from which coherence matrices were directly computed.

#### 4.3.1 Motif enumeration and structural data integration

To evaluate the local regulatory architecture within the empirical GRNs, we enumerated all non-isomorphic 2-node and 3-node directed, signed motifs using the dotmotif Python library. Statistical enrichment for each motif was assessed by comparing its observed frequency against a null distribution. This null background was generated by simulating ensembles of 50 independent randomized networks for each organism (see Supplementary Methods). Motif enrichment or depletion was subsequently quantified using empirical p-values derived from these shuffled ensembles. While exhaustive motif enumeration is efficient on sparse adjacency matrices, applying it directly to coherence matrices is computationally challenging. The integration of indirect paths generates a dense graph that overwhelms subgraph isomorphism algorithms. Consequently, we restricted motif enumeration to the adjacency matrix and employed transition-fate analysis to track how these localized topologies were structurally preserved or altered when mapped onto the continuous coherence matrix. To contextualize local architectures within the global hierarchy, constituent nodes of each enumerated motif were annotated with their hierarchical layer, functional NDA module, team identity, and transcription factor status. This integration enabled us to systematically map the precise functional and hierarchical localization of motifs across the networks.

### 4.4 iModulon analysis

The gene weight matrix *M* was obtained from the iModulon database, where columns represent independent component analysis modules, and rows represent genes [27]. Genes were included in a module if their absolute weight exceeded a curated threshold (default 0.05). Identifiers were mapped to standardized gene names, retaining locus IDs where necessary. The coherence matrix for *E. coli* was derived from the Abasy Atlas network and restricted to genes present in both datasets. Modules with fewer than two overlapping genes were excluded, and self-interactions were removed. For each unique gene pair (*i, j*), we computed the weight product *P*_*ij*_ = *w*_*i*_*w*_*j*_ as a signed measure of concordant or antagonistic contribution. Corresponding directional coherence values *C*_*ij*_ and *C*_*ji*_ were extracted. Pairs lacking both values were excluded. To establish the empirical regulatory sign, a consensus rule was applied: pairs were classified as cooperative if *C*_*ij*_ and *C*_*ji*_ shared the same sign (or if a single measured direction was positive), and antagonistic if their signs mismatched (or if a single measured direction was negative). For continuous magnitude comparisons, a composite coherence value was computed as the mean of *C*_*ij*_ and *C*_*ji*_ when both were present, or as the single available value otherwise. To assess relative interaction strength, we computed the Spearman rank correlation between *P*_*ij*_ and this composite coherence value per module. For classification performance, predictions based on the sign of *P*_*ij*_ (*P*_*ij*_ *>* 0 predicting cooperation) were evaluated against the consensus regulatory sign using precision, recall, and F1 scores. Analysis was restricted to transcriptionally active iModulons in wild-type *E. coli* MG1655; samples from adaptive evolution experiments were excluded to avoid genomic discrepancies with the reference regulatory network.

To investigate the relationship between static network topology and dynamic gene expression variance under global machinery perturbations, we downloaded and analyzed pre-processed public RNA-sequencing data for *Escherichia coli* K-12 MG1655 strains (GEO accession GSE178281). The dataset consists of cultures grown in varying concentrations of LB media (0.25×, 0.50×, and 0.75×), which induces a dose-dependent decrease in RNA polymerase (RNAP) concentration and triggers global perturbations in the transcriptional machinery [44]. For each condition, we filtered for significantly differentially expressed genes (adjusted p-value *<* 0.05) and compared their absolute log_2_ fold changes (absolute log_2_FC) against their corresponding absolute incoming coherence values to assess potential correlations between topological consistency and expression magnitude.

## 4.5 Acknowledgments

During the preparation of this work, the authors used Google Gemini to improve the manuscript’s readability. After using this tool, the authors reviewed and edited the content as needed and take full responsibility for the publication’s final content.

## 4.8 Author contributions

M.K.J. and P.H. designed research; M.K.J. and C.K. supervised research; P.H., R.J., and A.S.R. analyzed data; P.H. performed research and wrote the first draft; all authors contributed to editing the manuscript.

## 4.9 Data Availability

All code and data used in this study are publicly available in the GitHub repository: https://github.com/MoltenEcdysone09/StructuralCoherence. This repository includes all analysis scripts, plotting code, and input datasets required to reproduce the results presented in this manuscript.

## Appendices

### A Supplementary Methods

#### A.1 Teams Identification Algorithm

Team identification proceeded through a multi-stage clustering and reassignment algorithm applied to the coherence matrix, *C*. To isolate the underlying regulatory logic, continuous coherence values were first discretized using the sign function, yielding a matrix *S* where *S*_*ij*_ = sgn(*C*_*ij*_) ∈ {−1, 0, 1}. Because teams intrinsically require mutual activation, any nodes lacking strictly positive self-activation (*S*_*ii*_ ≤ 0) were immediately isolated as permanent singleton teams and excluded from further merging.

For the remaining nodes, we established initial core groups using complete-linkage hierarchical clustering. Nodes were clustered together only if their bidirectional interactions were strictly positive (min(*S*_*ij*_, *S*_*ji*_) *>* 0) or mutually undefined (NaN). This strict linkage ensures that initial groups contained no conflicting or neutral regulatory signals. These core groups were subsequently split into connected components using undirected positive interactions to guarantee topological contiguity.

The algorithm then refined these initial boundaries using a sequential greedy node-migration process. Nodes were permitted to migrate to larger or equal-sized groups, provided the move introduced no negative interactions. In the first optimization stage, node migrations were greedily executed to maximize the number of positive intra-team connections gained. Once these moves were exhausted, a second stage evaluated migrations to maximize the number of negative interactions segregated outside the target team.

Finally, for densely connected Erdős–Rényi (ER) networks, a concluding connected-component split was applied to guarantee that all final teams formed fully contiguous positive subnetworks (Algorithm 1). However, this final spatial splitting was deliberately omitted for artificial hierarchical (HI) networks and whole-organism GRNs. In these inherently sparse structures, the vast majority of nodes are terminal input or output genes that can propagate or receive inhibitory signals without participating in mutual positive cycles. Enforcing strict positive contiguity on these peripheral nodes artificially fragments them into isolated singletons, drastically inflating the total team count without reflecting actual underlying regulatory conflict.

We also note that the greedy nature of the refinement phase has boundary cases in which a node satisfies the compatibility and connectivity criteria for multiple teams simultaneously. In such situations, the final assignment of that node may depend on the order in which eligible teams are evaluated. Importantly, while this ambiguity can affect the specific membership of individual teams, it does not alter the procedure’s outcome with respect to the total number of teams identified, ensuring the robustness of all reported results to these ordering-induced degeneracies.

#### A.2 Generation of artificial networks

The base topologies for our synthetic GRNs were derived from an exhaustive enumeration of all unique, non-isomorphic directed signed graphs of two and three nodes. Triad configurations were generated using the signed triad census formalism, yielding 138 unique topologies. To complete the structural ensemble, we explicitly enumerated all 6 unique directed signed dyads, bringing the total comprehensive library to 144 unique base regulatory circuits. To systematically select circuits for the scaling analysis, we applied strict structural filters to this ensemble. First, we discarded trivially disconnected graphs. Crucially, we required that every selected circuit possess at least one “middle node”—defined topologically as a node with at least one incoming and one outgoing edge. This constraint ensures that the selected circuits are inherently capable of signal integration and propagation, a strict prerequisite for circuits functioning within the regulatory core of larger biological networks. Furthermore, we excluded circuits with mathematically undefined baseline structural coherence. Such circuits (e.g., the 010-N0 dyad, which consists of only a single inhibitory edge) lack the cyclic or reinforcing pathways necessary to form mutually activating functional teams upon scaling, rendering them trivial for an analysis of coordination.

##### Algorithm 1

Team Identification via Coherence Matrix Partitioning and Vectorized Refinement

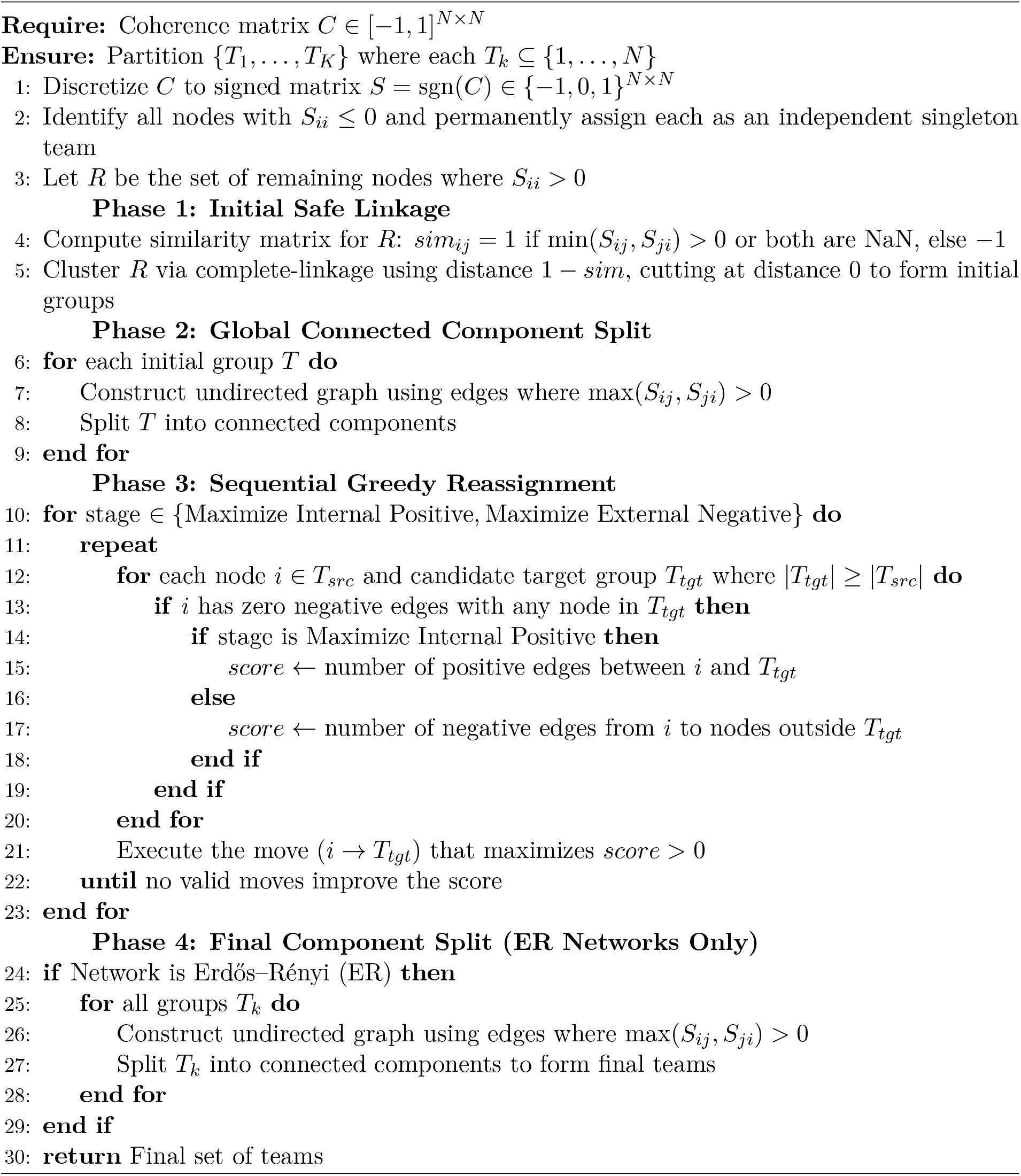

The functionally viable circuits were then categorized into fundamental topological classes based on their MAN (Mutual, Asymmetric, Null) nomenclature: feed-forward (030T), cyclic (030C), and complete (100 for dyads and 300 for triads) (Figure S1). To ensure our scaled ensembles spanned the full spectrum of structural coherence without redundant computation, we implemented a targeted sampling algorithm. For each topological class, we first isolated the No Self-Activation (NS) variants and evaluated their baseline structural coherence. To prevent sampling topologically distinct but dynamically redundant architectures, we filtered out circuits with duplicate structural coherence values (applying a numerical tolerance of 0.1). From the remaining unique circuits within each class, we sampled up to four representative circuits corresponding to the maximum (best), minimum (worst), 25th percentile, and 75th percentile of the structural coherence distribution.

These selected representative circuits were formally designated using their full MAN codes. The MAN code prefix indicates the absolute count of edge types within the circuit (e.g., “300” denotes 3 mutual, 0 asymmetric, and 0 null connections). This is followed by an edge-composition string (e.g., “NNNPPP” for triads, or “PN” for dyads) that specifies the exact sequence of activating (P), inhibitory (N), or absent (0) interactions. This string maps sequentially to the specific directed edges of the base circuit: *A* → *B* and *B* → *A* for dyads, and *A* → *B, B* → *A, A* → *C, C* → *A, B* → *C*, and *C* → *B* for triads.

Each selected base circuit was expanded into a larger network using a block-expansion procedure. For a circuit of size *M* ∈ {2, 3}, each node was expanded into a coordinated group of *S* nodes, utilizing scaling factors *S* ∈ {10, 30, 50}. This yielded unsparsified baseline networks of total size *N* = *M* · *S*. To assess the role of localized auto-regulation, we evaluated two conditions: Self-Activation (SA) and No Self-Activation (NS). In the SA condition, the diagonal elements of the base adjacency matrix *A*_*base*_ ∈ {−1, 0, 1}^*M*×*M*^ were explicitly set to 1; in the NS condition, they remained 0. If *J*_*S*_ is an *S* × *S* matrix of ones, the adjacency matrix of the expanded network is given by

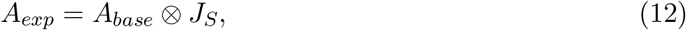

where ⊗ denotes the Kronecker product. Each entry (*A*_*base*_)_*ij*_ generates an *S* × *S* block in *A*_*exp*_ whose entries all carry the same sign, ensuring that the regulatory logic of the base circuit is inherited by the corresponding node groups.

We generated nested ensembles of Erdős–Rényi (ER) networks by varying the global edge density *ρ* ∈ [0.10, 1.00] in increments of 0.10. For a targeted density *ρ*, the number of directed edges *E* in the scaled network was constrained to *E* = *ρN* ^2^. For each combination of base circuit, scaling factor, and target density, we generated 50 independent replicates (except for the 100% density configuration, which has only one unique topology).

Networks of varying global densities (10% to 100%) were generated independently for each replicate rather than as strict additive subsets. However, to isolate the effects of density from random topological variance, each replicate was constructed around a deterministic structural invariant. For any given network size and replicate seed, the generation algorithm first extracted a weakly connected spanning forest from the allowed circuit-induced blocks. Because the random state was conserved per replicate, this initial spanning backbone was exactly identical across all densities of that replicate, acting as a shared, immutable scaffold. Following the placement of this conserved backbone, the additional edges required to reach the target global density were populated stochastically. Consequently, across all densities of the same base topological connectivity (cyclic, feed-forward and complete), network size, and replicate, the fundamental connectivity structure is perfectly conserved, differing only in the randomly distributed auxiliary edges.

##### A.2.1 Generation of artificial hierarchical networks

Artificial hierarchical (HI) networks were generated using the same block-expansion and edge-growth framework, with additional constraints to enforce biologically motivated hierarchy. Feed-forward circuits (030T) were excluded from this process, as embedding an inherently hierarchical circuit into a macro-hierarchical structure is topologically redundant. Nodes were assigned to three hierarchical layers—input, middle, and output—using fixed proportions derived from empirical GRNs (5% Input, 5% Middle, 90% Output). Structural organization was enforced via layer-wise block masks defining admissible edge placements. Connectivity followed a feed-forward architecture: edges were permitted from the input to the middle and from the middle to the output layers. Intra-layer connections were prohibited in the input and output layers but allowed in the middle layer.

Block-wise densities were fixed to mirror empirical coupling strengths: 100% for Middle-to-Middle, 85% for Middle-to-Output, and 50% for Input-to-Middle. Direct Input-to-Output connectivity served as a control parameter, varied from 10% to 100% in 10% increments. During placement, priority was given to upstream-to-downstream connections to ensure downstream nodes received at least one incoming edge from an appropriate upstream block. Remaining edges within admissible blocks were placed uniformly at random until target densities were reached.

Under these constraints, the total number of edges *E*_total_ in a hierarchical network of scale *S* can be written as a function of *S* and the variable input-to-output density *D*:

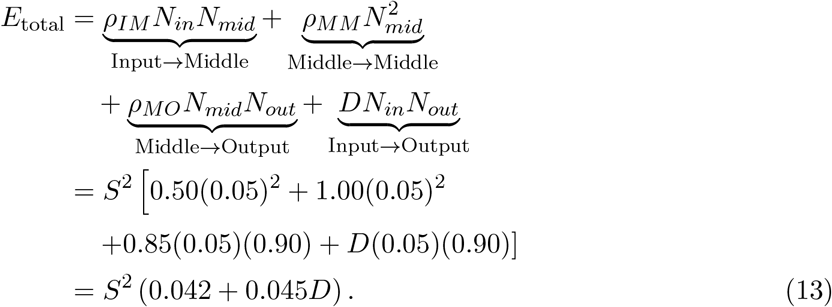

Here, *N*_*in*_ = *N*_*mid*_ = 0.05*S* and *N*_*out*_ = 0.90*S* represent the node counts per layer. The block connection densities are fixed at *ρ*_*IM*_ = 0.50, *ρ*_*MM*_ = 1.00, and *ρ*_*MO*_ = 0.85. The constant term (0.042) thus aggregates the edges from these fixed-density blocks, while the variable term (0.045*D*) accounts for the variable Input-to-Output connectivity. Since the total number of nodes is *N*_total_ = *S*, a fully connected unconstrained network with self-loops would contain *S*^2^ edges; thus, the imposed hierarchical constraints limited the network to a maximum of approximately 8.7% of its theoretical full connectivity.

Following edge placement at each density level, a minimal connectivity check was applied to ensure that every node participated in at least one interaction within its assigned team. For every node in a downstream layer (middle or output), the algorithm verified the presence of at least one incoming directed edge from an upstream node belonging to the same expanded base node (i.e., the identical structural team). If no such intra-team connection existed, a valid upstream intra-team edge was randomly selected and added. After satisfying these baseline global connectivity and lineage constraints, the remaining edge budget for each hierarchical block was populated uniformly at random until the specific target block densities were exactly met. For each targeted density level, 50 independent replicates were generated.

#### A.3 Network connectivity and functional enrichment analyses of whole-organism GRNs

##### A.3.1 Node-level coherence measures and topological classification

To characterize regulatory roles, we computed two scalar quantities for each gene *i* from the coherence matrix *C*: absolute outgoing coherence (Σ_*j*_ |*C*_*ij*_|) and absolute incoming coherence (Σ_*j*_ |*C*_*ji*_|). Based on these measures, nodes were classified into three categories: input nodes, defined by nonzero outgoing and zero (or undefined) incoming coherence; output nodes, defined by nonzero incoming and zero (or undefined) outgoing coherence; and middle nodes, which exhibit both incoming and outgoing coherence. This classification is structurally equivalent to a standard adjacency-based topological definition, wherein input nodes possess an in-degree of zero, output nodes possess an out-degree of zero, and middle nodes possess both nonzero in- and out-degrees. We compute these classifications directly from the coherence matrix to maintain analytical consistency with the coherence framework utilized throughout the study.

##### A.3.2 Generation of randomized whole organism GRNs

To establish a topological baseline, we generated 50 randomized null models for each wild-type GRN using a degree- and sign-preserving edge-swapping algorithm. By iteratively swapping target nodes of same-sign edges, this method shuffles the wiring of the GRN. Consequently, each randomized network maintains the exact nodal degrees and the global regulatory-sign distribution, ensuring that observed structural deviations reflect higher-order topological organization rather than fundamental connectivity constraints.

##### A.3.3 Inter and Intra-layer connectivity and density analysis

To quantify connectivity patterns between topological layers, we computed edge densities between all pairs of node classes (input, middle, output). For a directed layer pair (*α* → *β*), the density was defined as

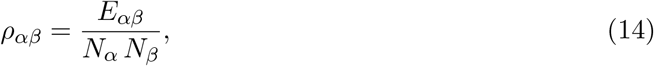

where *E*_*αβ*_ denotes the number of observed directed edges from nodes in layer *α* to nodes in layer *β*, and *N*_*α*_ and *N*_*β*_ denote the number of nodes in the source and target layers, respectively. This normalization corresponds to the maximum number of possible directed edges consistent with both directionality and layer membership.

##### A.3.4 Module annotations and intra-versus inter-module connectivity

Functional module annotations were obtained directly from Abasy Atlas, which employs the Natural Decomposition Approach (NDA) to partition GRNs into global regulators, basal cellular machinery, autonomous functional modules, and inter-modular genes [42]. For each network, module sizes were quantified as the number of genes assigned to each NDA component. To analyze coupling within and between modules, regulatory interactions were classified as intra-module if both genes belonged to the same module and inter-module otherwise. Densities of intra- and inter-module connections were then computed using the same formula as described for the inter and intra-layer connectivity analysis, and further stratified by the coherence-defined topological layers of the interacting genes.

##### A.3.5 Functional annotation and enrichment analysis

Gene Ontology (GO) [55, 56] annotations were retrieved from UniProt [57] using the bioservices [58] and goatools [59] Python libraries. Treating the Molecular Function GO as a directed acyclic graph (DAG), we assigned genes to categories via ancestral traversal, retaining only terms with a minimum DAG depth of 5 to reduce redundancy. Genes mapping to the ancestry of “transcription regulator activity” (GO:0140110) were classified as transcription factors, ensuring consistent identification despite heterogeneous annotation depths.

To evaluate associations between topological roles (Input, Middle, Output) and either specific biological functions or predefined Abasy Atlas NDA modules, we performed enrichment analyses. We utilized a one-sided Fisher’s Exact Test comparing target role membership against the entire network, followed by Benjamini–Hochberg false discovery rate (BH-FDR) correction (*α* = 0.05).

Significant GO terms were ranked using a Combined Score that integrates both enrichment magnitude and term specificity:

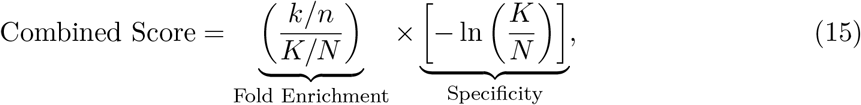

where *k* and *n* represent the number of annotated genes and the total genes within the specific topological role, respectively, while *K* and *N* denote the corresponding values for the global network population. This formulation systematically upweights highly specific, over-represented terms while penalizing broad background annotations.

### B Supplementary Figures

**Figure S1:**
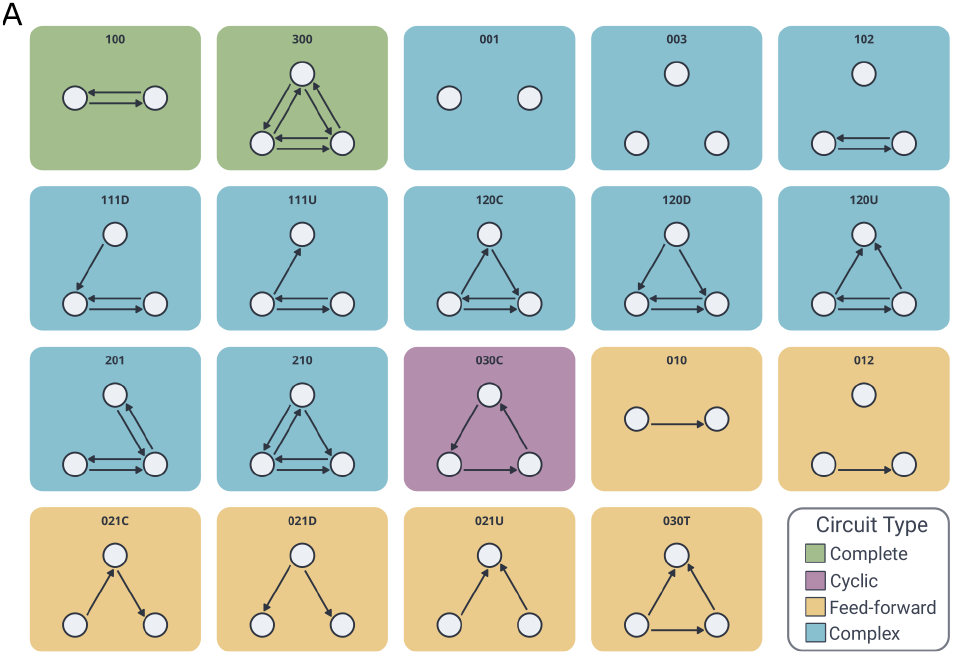
Topological classification of unique dyadic and triadic network circuits. **(A)** Wiring diagrams of directed 2-node and 3-node network circuits, labeled by their corresponding Mutual Asymmetric Null (MAN) naming convention. The structural topologies are stratified into four distinct classes, indicated by background shading, highlighting varying degrees of hierarchical organization and structural coherence within the network space. Complete (Green): Fully connected subgraphs characterized exclusively by bidirectional interactions between all constituent nodes. Cyclic (Purple): Architecture containing strictly directed, closed-loop feedback. Feed-forward (Yellow): Directed acyclic topologies. This category includes canonical feed-forward loops (030T), linear cascades (021C), diverging/converging cascades (021D, 021U), and asymmetric dyads (010). Complex (Blue): Intermediate and composite structures featuring a mixture of asymmetric, mutual, and null edge pairs (e.g., 111D, 120C, 210) that fall outside the strict acyclic or fully connected definitions of the other functional categories. Nodes denote individual regulatory elements, and arrows indicate directed connections. Missing edges represent the strict absence of an interaction. Additionally, the completely disconnected null circuits (001 and 003) lack any directed interactions and are classified as Complex, as they do not satisfy the structural criteria of the other categories.

**Figure S2:**
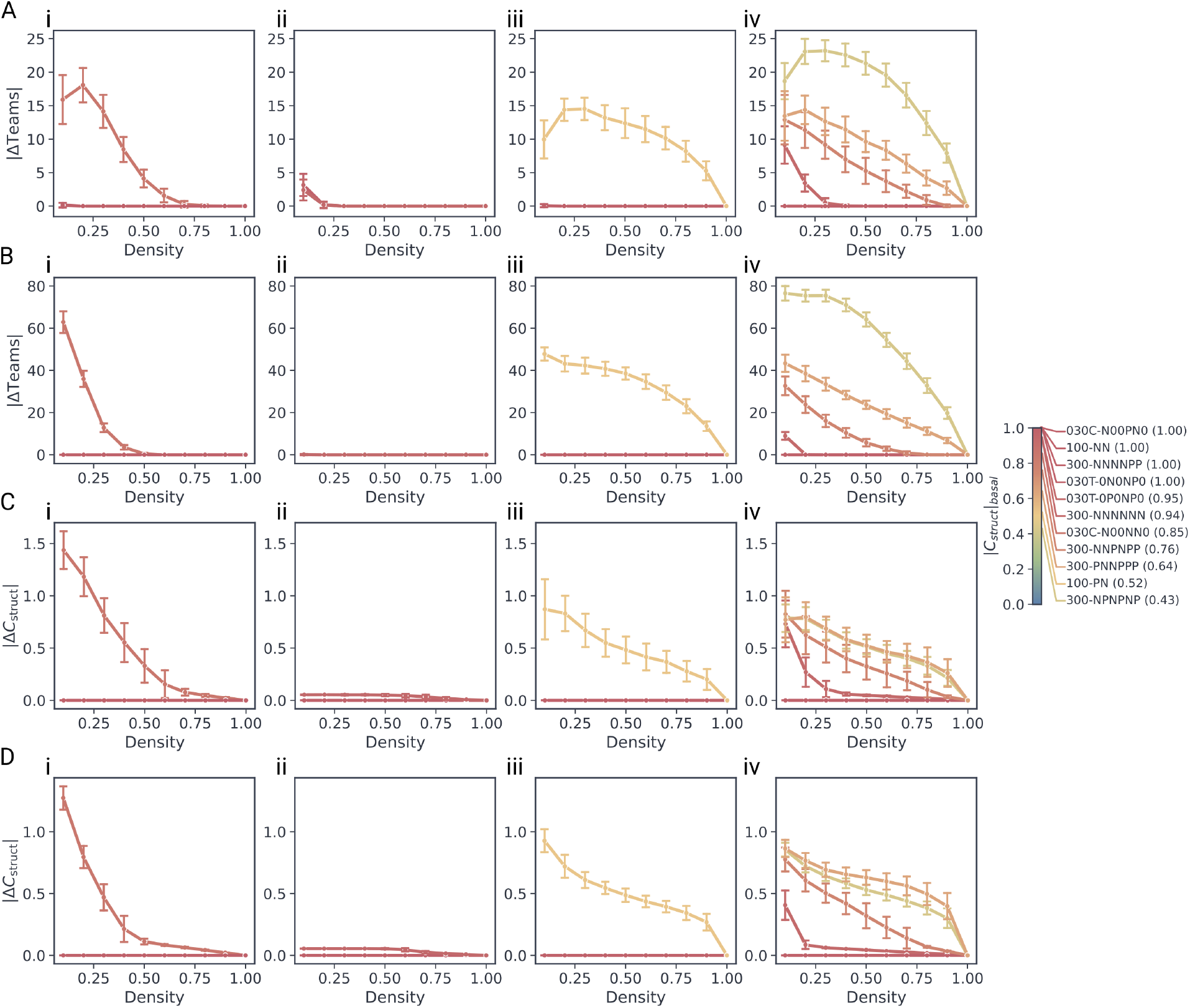
Absolute error trajectories for 10× and 30× scaled networks under sparsification. Line plots tracking the absolute error in team count (|ΔTeams|) for (**A**) 10× and (**B**) 30× scaled networks, and absolute error in structural coherence (|ΔC_struct_|) for (**C**) 10× and (**D**) 30× scaled networks as a function of global edge density (%). Columns segregate results by topological class: (**i**) cyclic, (**ii**) feed-forward, (**iii**) complete 2-node, and (**iv**) complete 3-node architectures. Error bands denote the standard deviation across 50 independent replicates.

**Figure S3:**
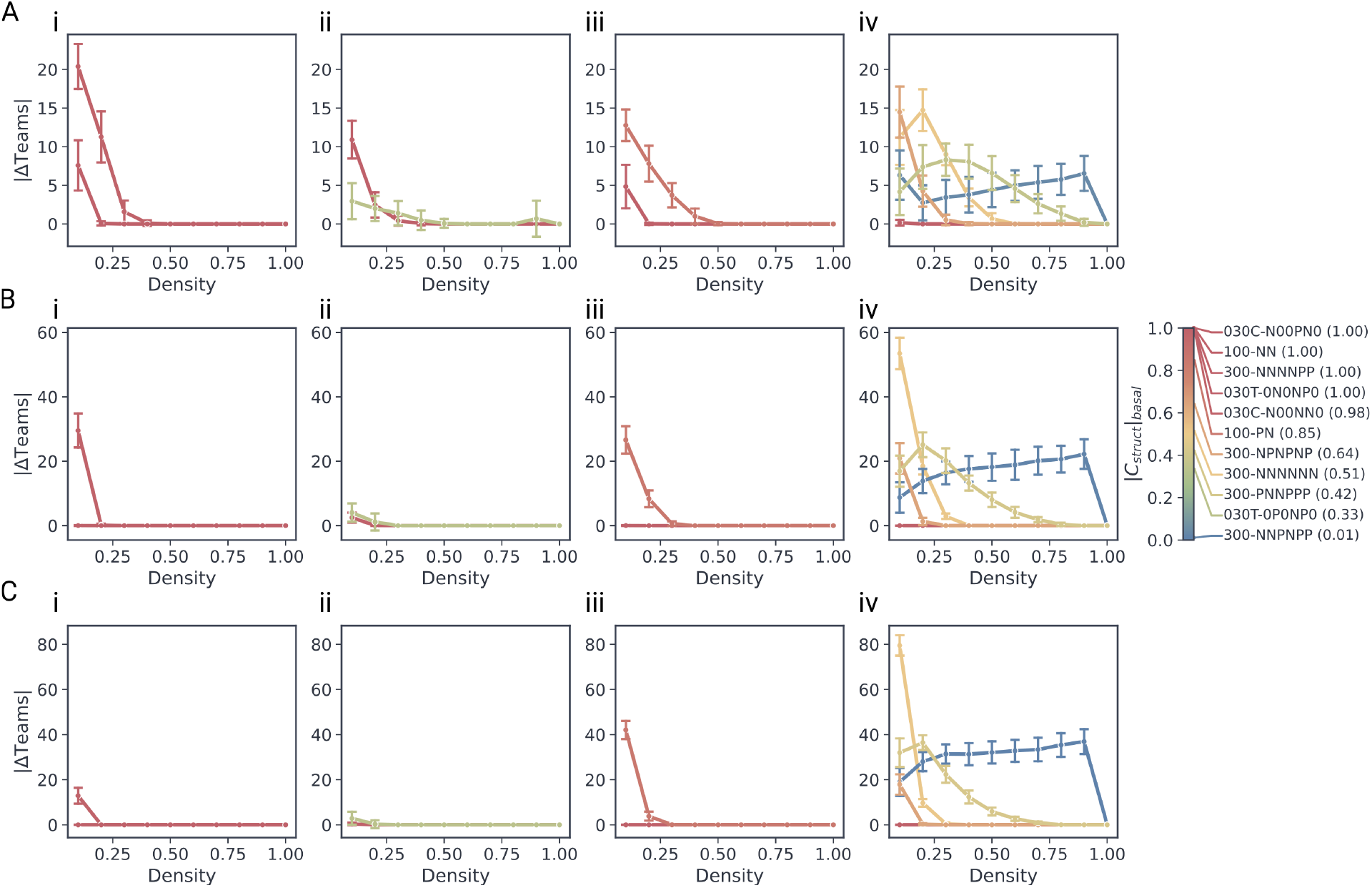
Absolute error trajectories in team count for No Self-Activation (NS) ensembles under sparsification. Line plots tracking the absolute error in team count (|ΔTeams|) for (**A**) 10×, (**B**) 30×, and (**C**) 50× scaled networks as a function of global edge density (%) within the NS ensemble. Columns segregate results by topological class: (**i**) cyclic, (**ii**) feed-forward, (**iii**) complete 2-node, and (**iv**) complete 3-node architectures. Error bands denote the standard deviation across 50 independent replicates.

**Figure S4:**
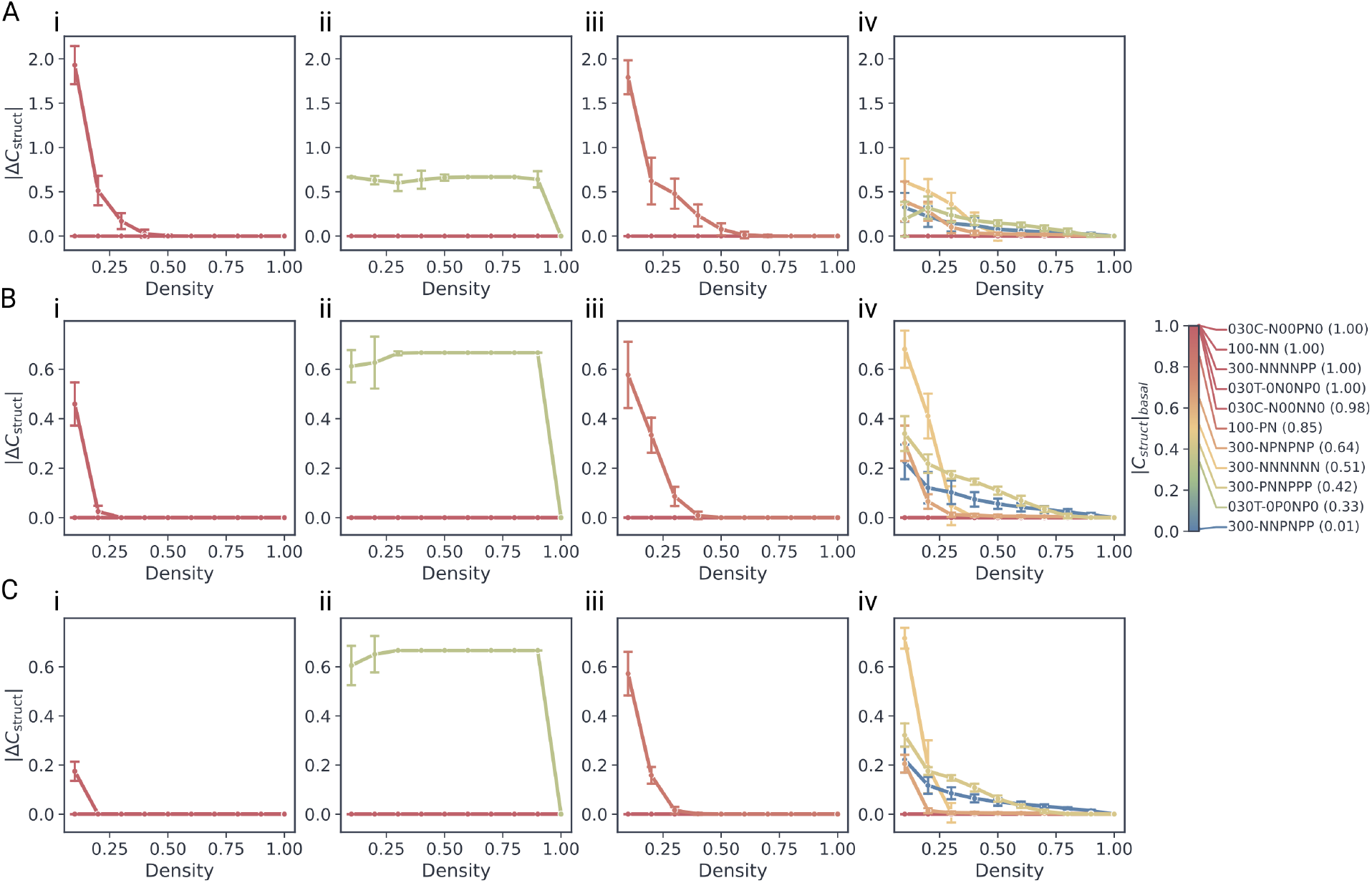
Absolute error trajectories in structural coherence for No Self-Activation ensembles (NS) under sparsification. Line plots tracking the absolute error in structural coherence (|ΔC_struct_|) for (**A**) 10×, (**B**) 30×, and (**C**) 50× scaled networks as a function of global edge density (%) within the NS ensemble. Columns segregate results by topological class: (**i**) cyclic, (**ii**) feed-forward, (**iii**) complete 2-node, and (**iv**) complete 3-node architectures. Error bands denote the standard deviation across 50 independent replicates.

**Figure S5:**
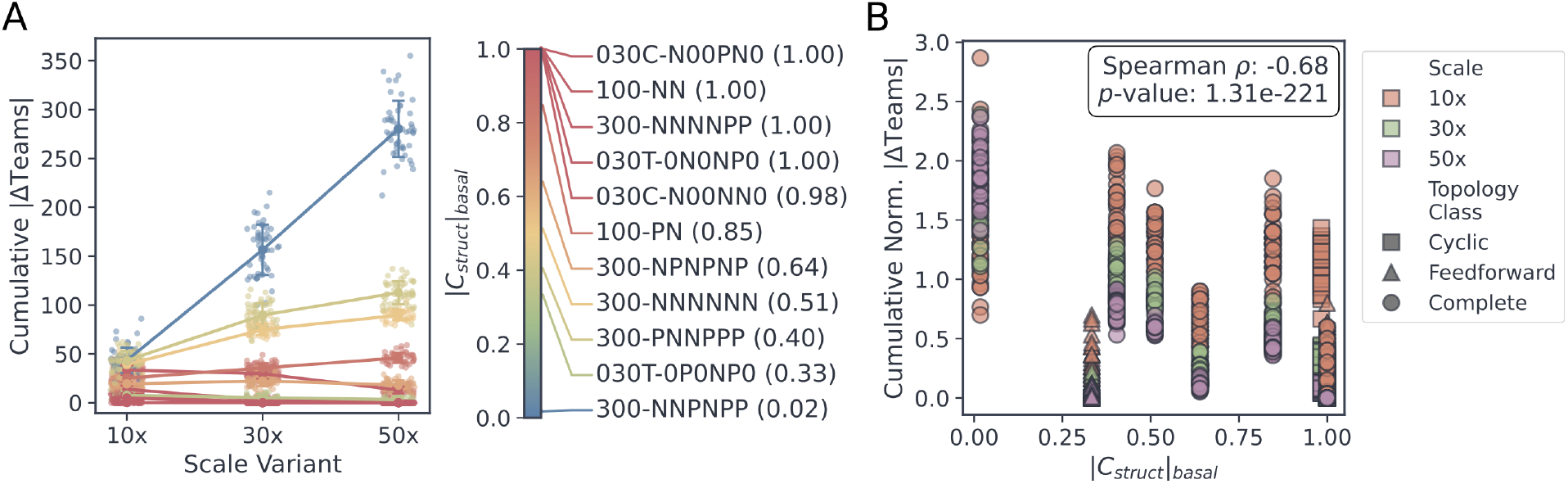
Cumulative error distributions and baseline coherence correlations for No Self-Activation ensembles. (**A**) Comparison of the distribution of Cumulative Absolute Error (CAE) across the scaling factors (10×, 30×, 50×) for all scaled NS circuits. (**B**) Scatter plot correlating Normalized Cumulative Absolute Error (Norm-CAE) against the absolute baseline structural coherence (|C_struct_|) for the NS networks. Individual markers represent unique circuit configurations across all scales and topology classes. The Spearman rank correlation coefficient (*ρ*) and associated p-value are indicated in the panel background.

**Figure S6:**
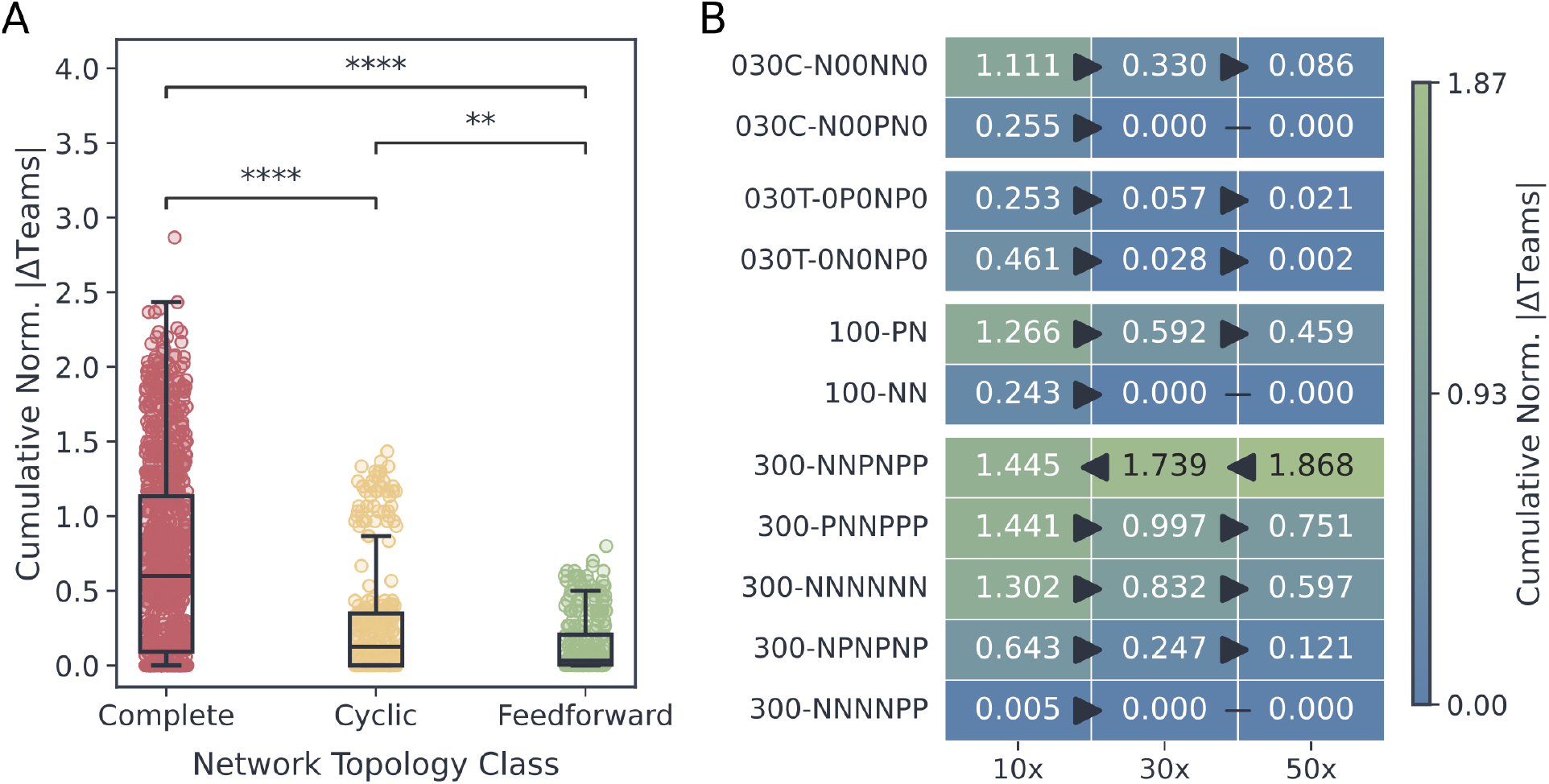
Topological resilience and scale-dependent buffering of NS networks under sparsification. (**A**) Box plots comparing the distribution of Normalized Cumulative Absolute Error (Norm-CAE) across topological classes for the NS networks. Statistical comparisons between distributions were performed using the Mann–Whitney U test; significance levels are denoted as ns (*p >* 0.05), * (*p* ≤ 0.05), ** (*p* ≤ 0.01), *** (*p* ≤ 0.001), and **** (*p* ≤ 0.0001). (**B**) Heatmap displaying the mean Norm-CAE values for each circuit across network scaling factors. Connecting lines between adjacent cells indicate statistical relationships; arrows point from a higher mean to a statistically significant lower mean, whereas flat lines without arrowheads denote no statistically significant difference using the Mann-Whitney U test.

**Figure S7:**
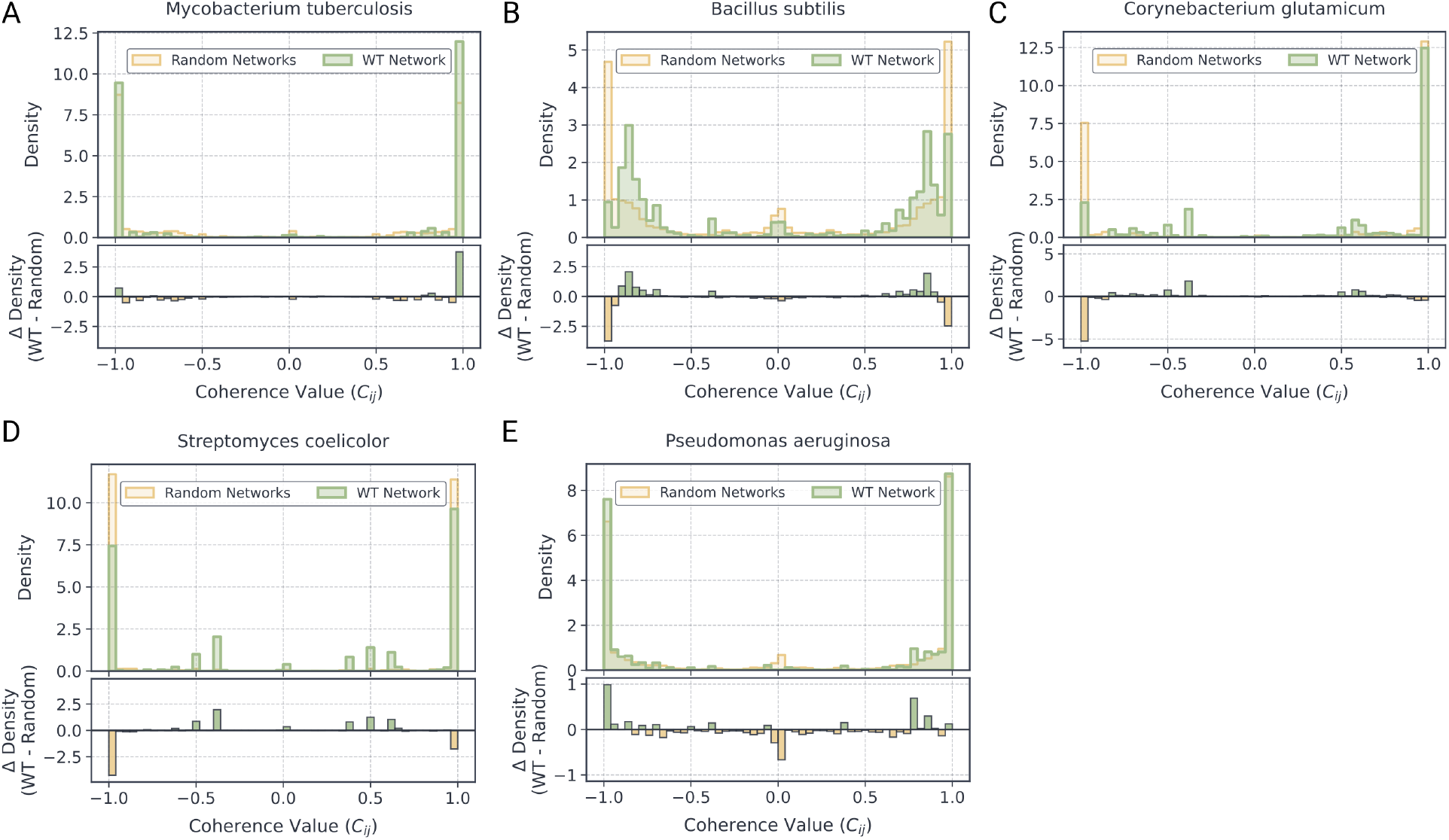
Organism-specific coherence distributions for WT and randomized networks. (**A-E**) Probability density distribution of coherence values comparing aggregated wild-type (WT) whole-organism GRNs against their degree-preserving randomized networks. The inset depicts the density difference (Δ Density = WT − Random).

**Figure S8:**
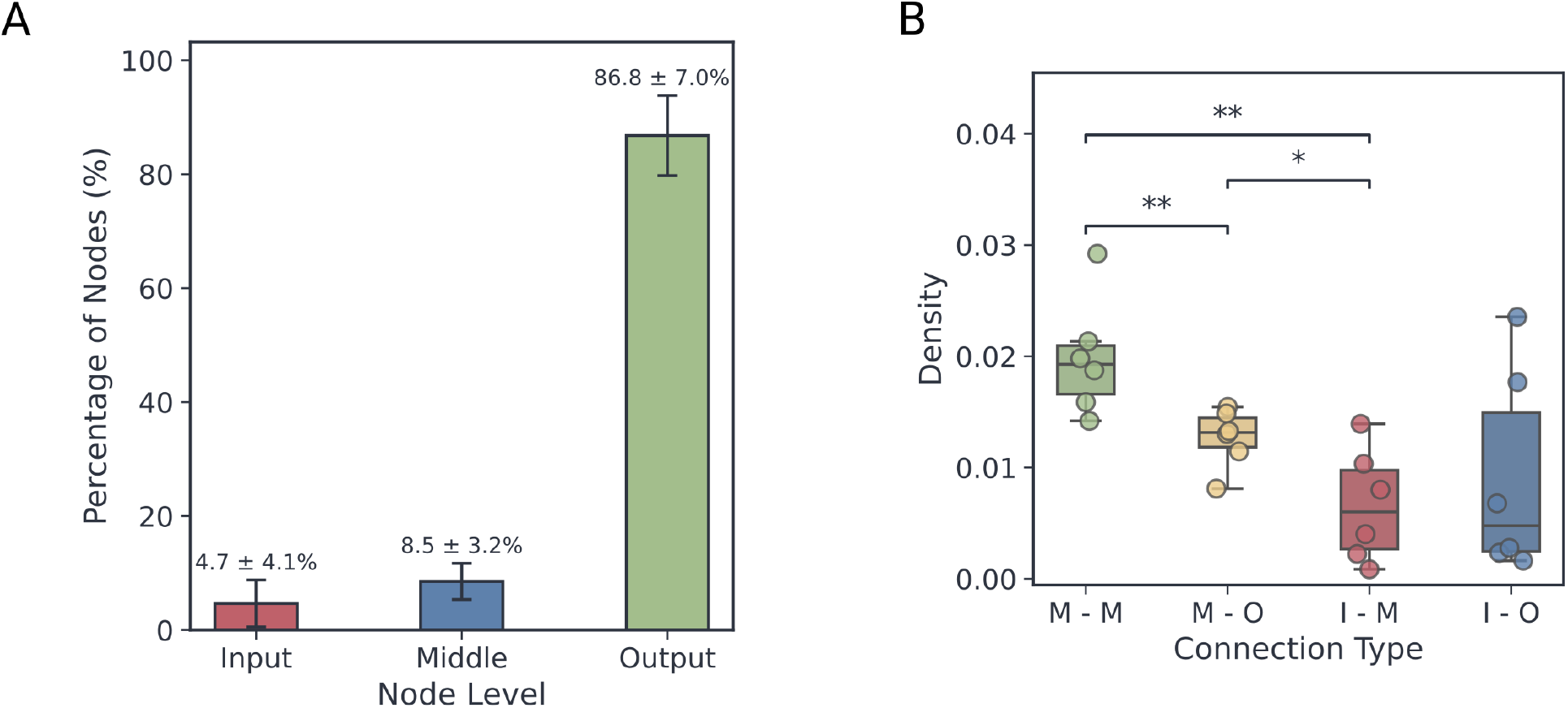
Hierarchical structure in whole-organism GRNs. (**A**) Bar plot detailing the percentage of Input, Middle, and Output genes across the whole-organism networks. (**B**) Box plots comparing inter-layer interaction densities (I: Input, M: Middle, O: Output, denoted as Source-Target) calculated from both directed adjacency matrices across all organisms. Statistical comparisons between distributions were performed using the Mann–Whitney U test; significance levels are denoted as ns (*p >* 0.05), * (*p* ≤ 0.05), ** (*p* ≤ 0.01), *** (*p* ≤ 0.001), and **** (*p* ≤ 0.0001).

**Figure S9:**
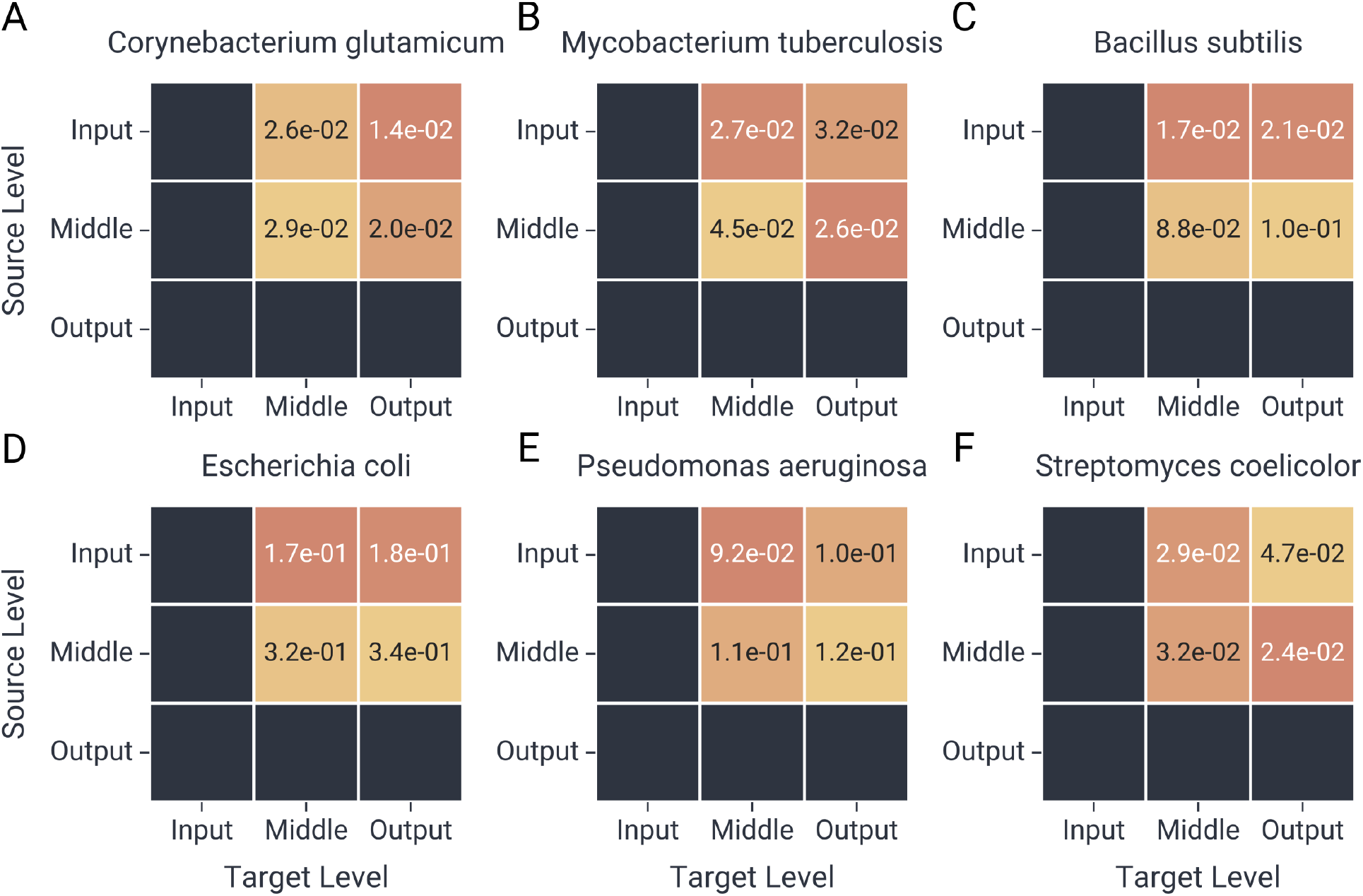
Organism-specific inter-layer interaction densities. (**A-E**) Heatmaps detailing the directed connection densities between the defined topological layers (Input, Middle, Output) derived from coherence matrices for each of the six analyzed organisms.

**Figure S10:**
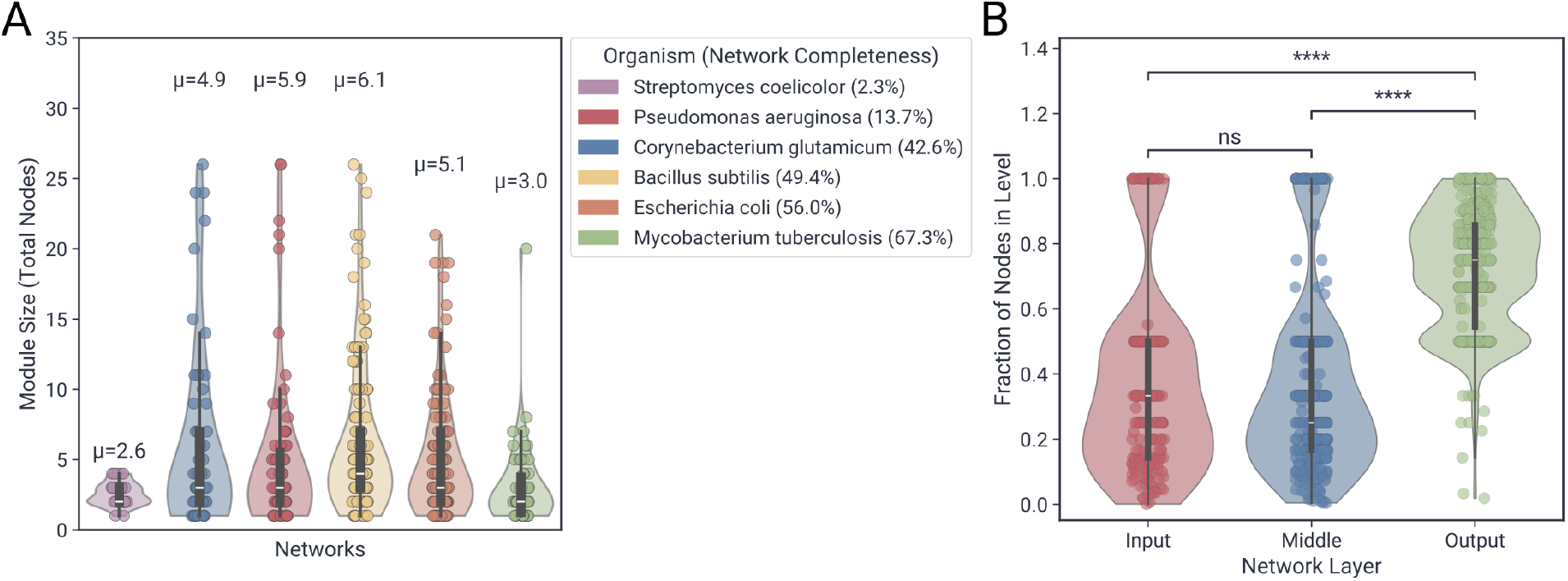
Structural composition and connectivity of biologically annotated functional modules. (**A**) Violin plots showing the distributions of functional module sizes (number of genes) across all six analyzed organisms, excluding modules above the 95th percentile. (**B**) Violin plots quantifying the fraction composition of Input, Middle, and Output genes within the defined functional modules.

**Figure S11:**
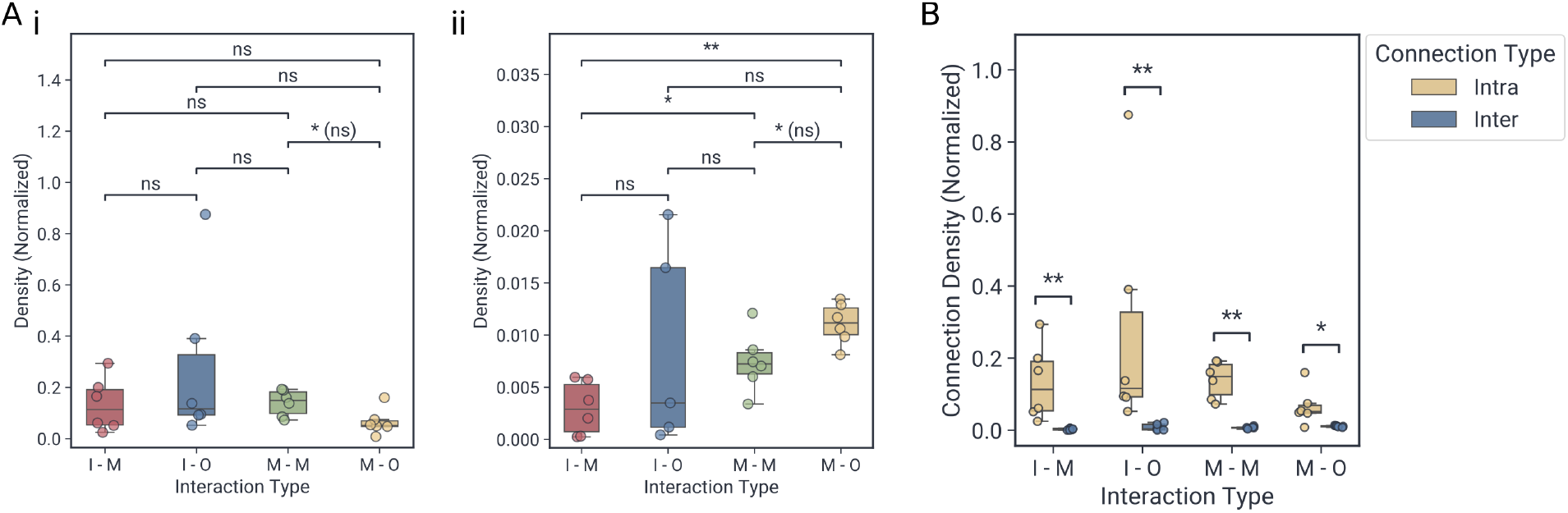
Layer-specific density distributions of intra- and inter-modular interactions. Box plots comparing the interaction densities between hierarchical layers (I: input, M: middle, O: output, denoted as source-target) separated into (**A**) inter-module and (**B**) intra-module connections. (**C**) Box plots comparing the interaction density between intra-module and inter-module for different layer connection combinations. Statistical comparisons between distributions were performed using the Mann–Whitney U test; significance levels are denoted as ns (*p >* 0.05), * (*p* ≤ 0.05), ** (*p* ≤ 0.01), *** (*p* ≤ 0.001), and **** (*p* ≤ 0.0001).

**Figure S12:**
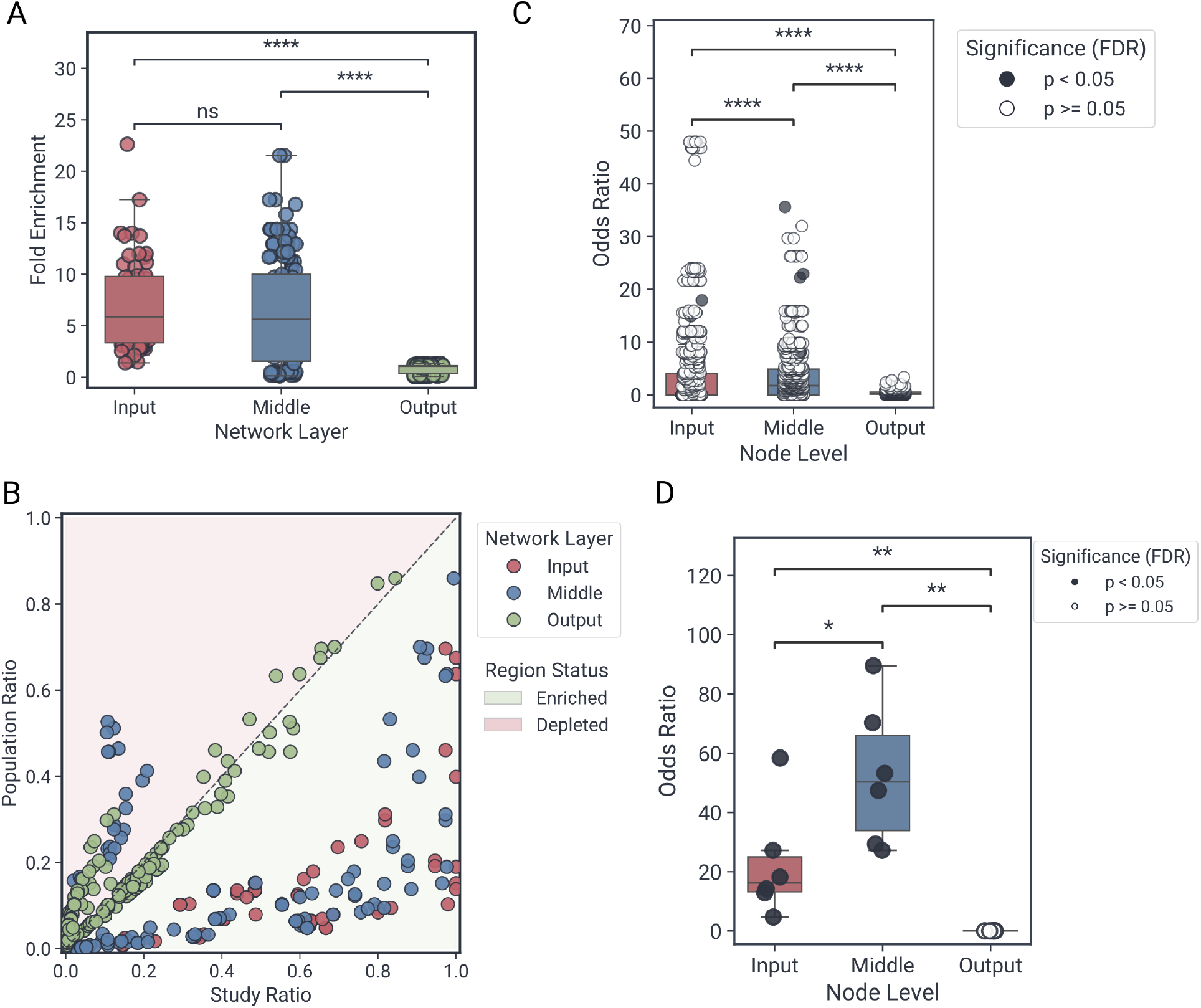
Global enrichment profiles and output layer characterization. (**A**) Box plots comparing the distributions of GO term enrichment odds ratios across the Input, Middle, and Output layers. (**B**) Scatter plot of layer-specific GO term gene ratios against background population ratio. The diagonal boundary delineates regions of functional enrichment (bottom right) from those of functional depletion (top left). (**C**) Box plots comparing the distributions of NDA module enrichment odds ratios across the three topological layers. (**D**) Box plots comparing the enrichment odds ratio specifically for the transcription regulator activity term (GO:0140110) across the three layers. Statistical comparisons between distributions were performed using the Mann–Whitney U test; significance levels are denoted as ns (*p >* 0.05), * (*p* ≤ 0.05), ** (*p* ≤ 0.01), *** (*p* ≤ 0.001), and **** (*p* ≤ 0.0001).

**Figure S13:**
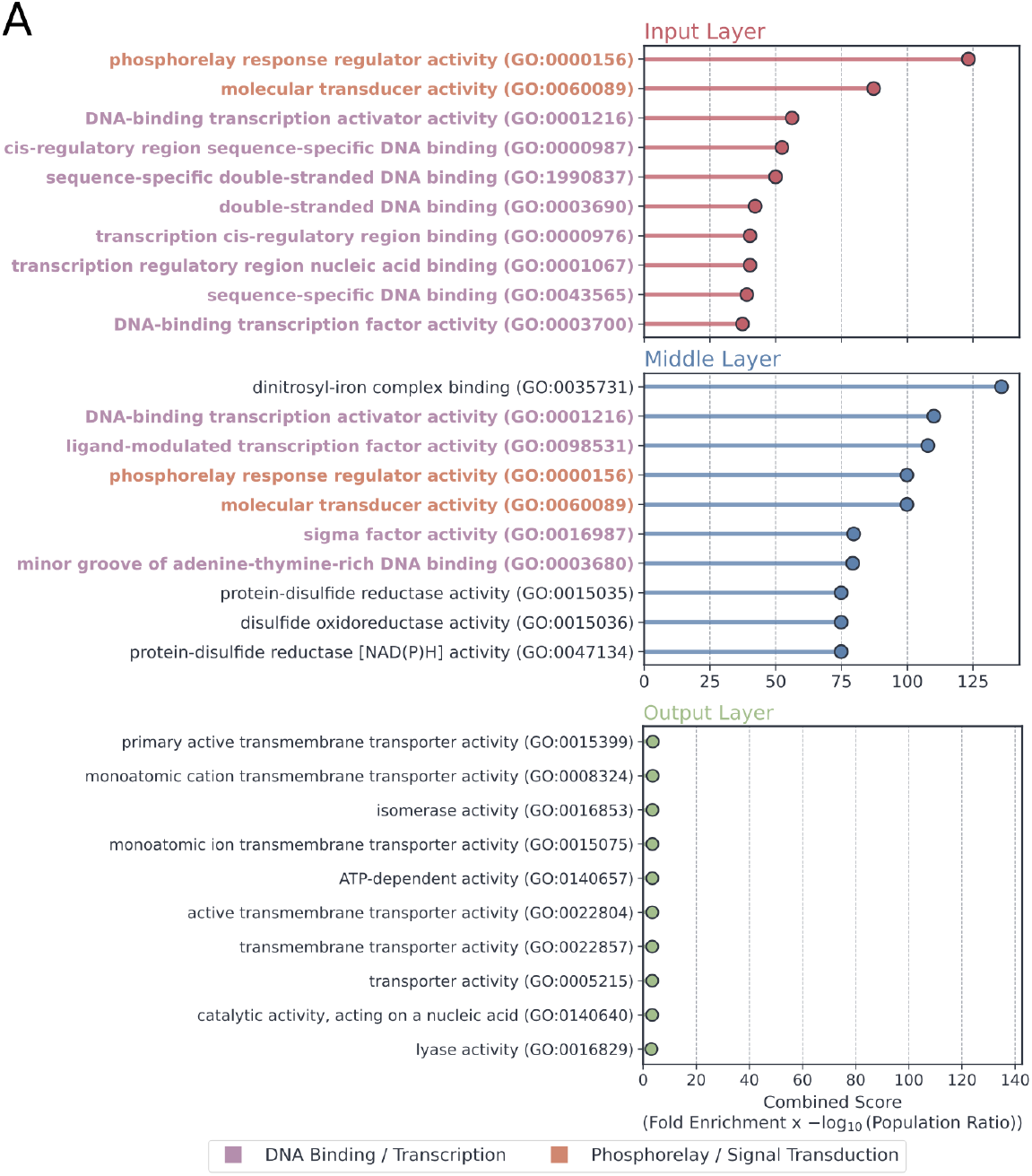
Functional specialization of the input and middle layer genes. **A** Lollipop plots detailing the top enriched Gene Ontology (GO) terms for the Input, Middle and Output layers across the analyzed whole-organism networks, ranked by their mean combined enrichment score across the six networks. Terms associated with DNA binding or transcription are highlighted in purple, while terms associated with phosphorelay or signal transduction are highlighted in orange.

**Figure S14:**
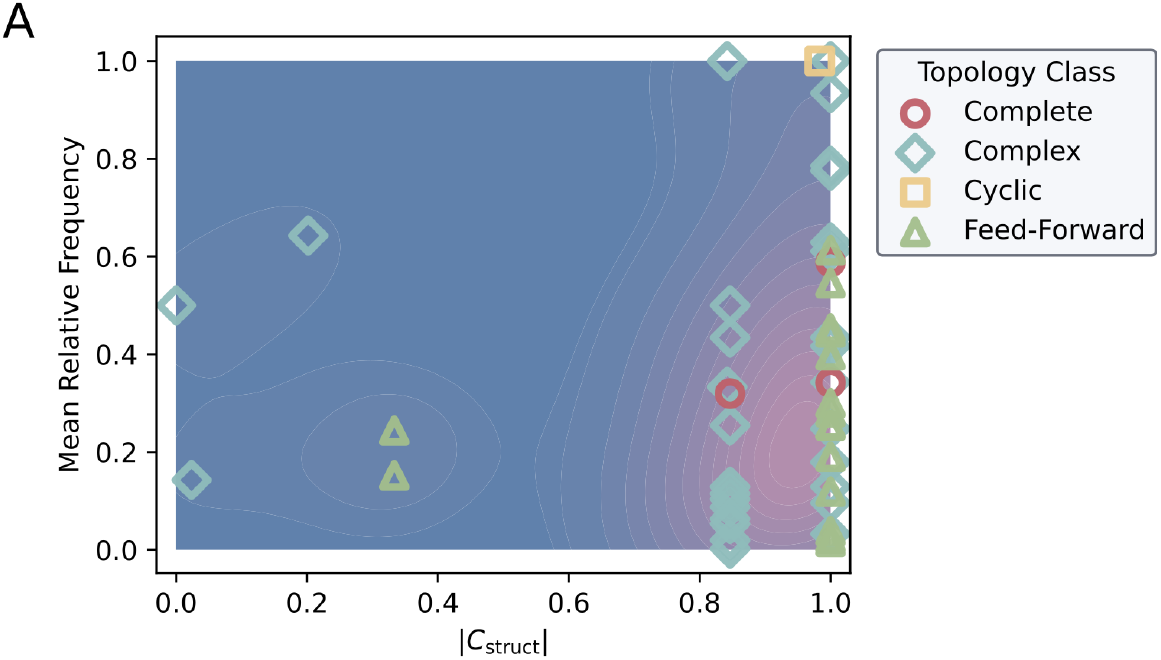
Relative frequency and structural coherence distribution of network motifs. (**A**) Scatter plot mapping the mean relative frequency of motifs across the organisms within their respective topological classes against their absolute structural coherence (|C_struct_|). Individual markers distinguish topological classes (feed-forward, cyclic, complete and complex) considered.

**Figure S15:**
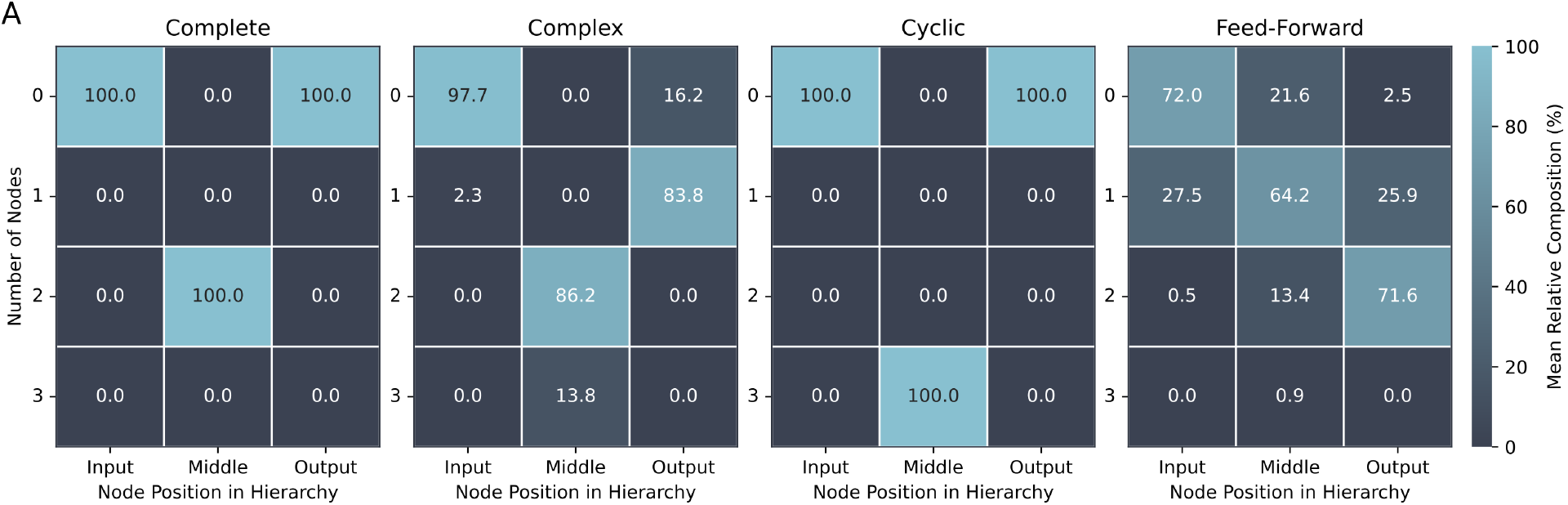
Distribution of hierarchical node counts across topological classes. Heatmaps displaying the mean percentage composition of the number of nodes situated in different hierarchical positions (input, middle, and output) across the four topological classes (feed-forward, cyclic, complete, and complex).

**Figure S16:**
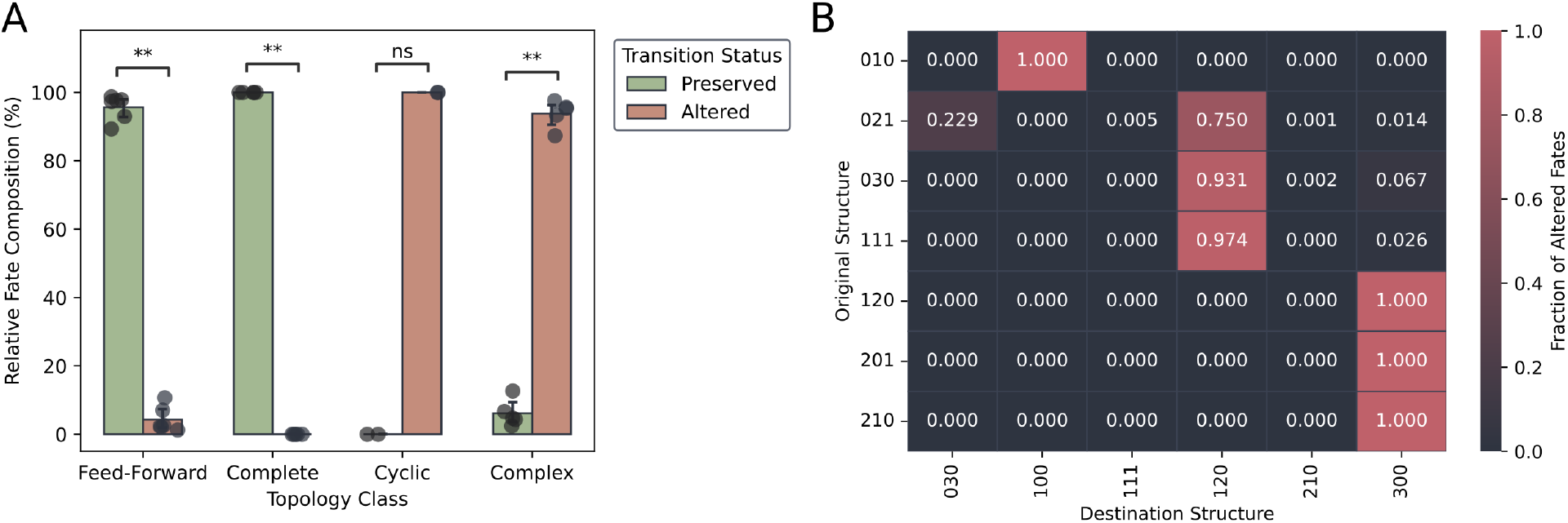
Topological transition rates and mapped pathways for altered motifs. (**A**) Bar plots with the fraction of structurally preserved versus altered motifs across the four topological classes. (**B**) Heatmap displaying the transition distribution of structurally altered motifs, mapping the original adjacency MAN code (y-axis) to the resulting coherence MAN code (x-axis).

**Figure S17:**
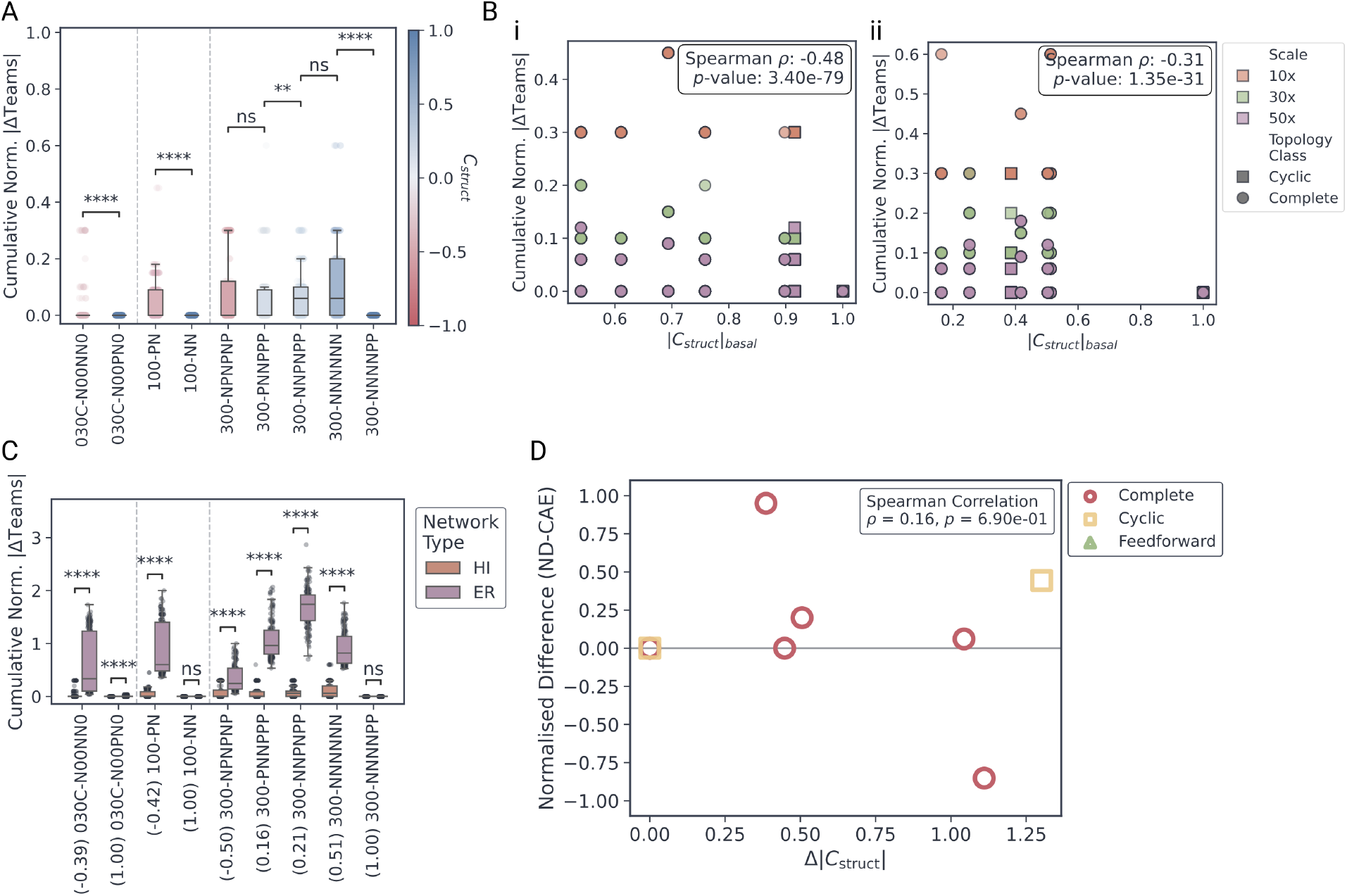
Structural robustness and coherence correlations for No Self-Activation hierarchical networks. (**A**) Box plots of the distribution of Norm-CAE across all scaling factors for individual base circuits within the NS hierarchical networks. Circuits are grouped by topological classes, separated by vertical dotted lines. Box colors map to the absolute structural coherence (|C_struct_|) of the corresponding base circuit. (**B**) Scatter plots correlating Norm-CAE against |C_struct_| for (**i**) SA and (**ii**) NS hierarchical (HI) networks. Individual markers represent unique circuit configurations across all scales and topological classes. The Spearman rank correlation coefficient (*ρ*) and associated p-value are indicated in each subpanel inset. (**C**) Box plots comparing Norm-CAE distributions between ER and HI NS networks for individual base circuits. X-axis labels denote the specific circuit MAN code and its baseline |C_struct_|. (**D**) Scatter plot correlating ND against the absolute structural coherence shift (Δ|C_struct_|) induced by self-activation for hierarchical networks. Individual markers differentiate topological classes. The Spearman rank correlation coefficient (*ρ*) and p-value are shown in the inset.

**Figure S18:**
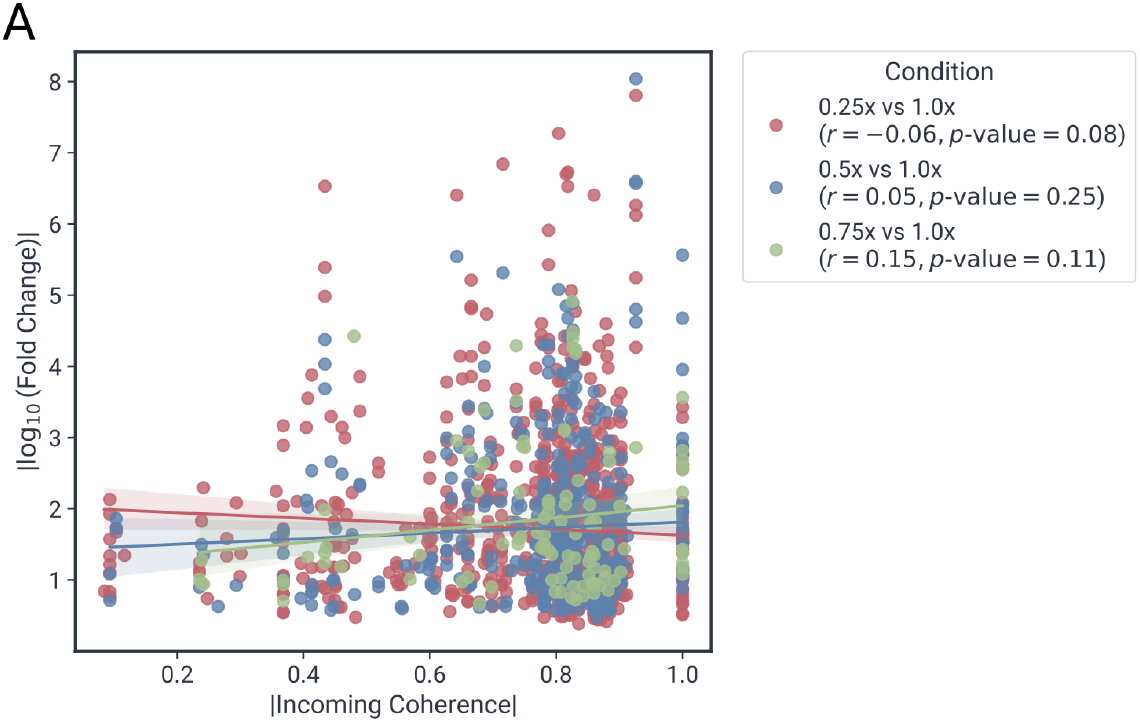
Analysis of gene expression variance under global transcriptional stress. **(A)** Scatter plots showing the relationship between absolute input coherence and absolute *log*_2_ fold change in gene expression measured in E. coli grown under progressively diluted LB media (0.25x, 0.50x, and 0.75x dilution).

